# High dimensional geometry of fitness landscapes identifies master regulators of evolution and the microbiome

**DOI:** 10.1101/2021.09.11.459926

**Authors:** Holger Eble, Michael Joswig, Lisa Lamberti, William B. Ludington

## Abstract

A longstanding goal of biology is to identify the key genes and species that critically impact evolution, ecology, and health. Yet biological interactions between genes (*1, 2*), species (*3–6*), and different environmental contexts (*7–9*) change the individual effects due to non-additive interactions, known as epistasis. In the fitness landscape concept, each gene/organism/environment is modeled as a separate biological dimension (*10*), yielding a high dimensional landscape, with epistasis adding local peaks and valleys to the landscape. Massive efforts have defined dense epistasis networks on a genome-wide scale (*2*), but these have mostly been limited to pairwise, or two-dimensional, interactions (*11*). Here we develop a new mathematical formalism that allows us to quantify interactions at high dimensionality in genetics and the microbiome. We then generate and also reanalyze combinatorically complete datasets (two genetic, two microbiome). In higher dimensions, we find that key genes (e.g. *pykF*) and species (e.g. *Lactobacillus plantarum*) distort the fitness landscape, changing the interactions for many other genes/species. These distortions can fracture a “smooth” landscape with one optimal fitness peak into a landscape with many local optima, regulating evolutionary or ecological diversification (*12*), which may explain how a probiotic bacterium can stabilize the gut microbiome.

## 1 Introduction

A fitness landscape depicts biological fitness as a function of its many underlying parts, namely genes, each as a separate dimension (*10, 13, 14*). Interactions between genes can change their individual impacts on fitness in a non-additive way (*15*), adding local peaks and valleys to the fitness landscape, which affects the evolutionary paths through the landscape (*16*). The mathematical frameworks to quantify biological interactions, namely epistasis, determine the degree of non-additivity, and the concept has been applied to genetics (*1, 2*), microbiomes (*3*), and ecology (*4–6*). In two dimensions, epistasis calculates interactions between e.g. two genes as the degree to which a double mutant phenotype can be predicted by measuring the two single mutants independently. Applying epistasis to genome-wide measurement of pairwise (*17, 18*) and three-way (*2*) genetic interactions has revealed biochemical pathways composed of discrete sets of genes as well as complex traits, such as human height, that are affected by almost every gene in the genome (*19, 20*). New techniques allow epistasis to be applied to broader data types (*21*).

Epistatic interactions can arise due to mutations (*13,14,22*) or when sex, recombination, and horizontal gene transfer bring groups of genes together (*1, 23–26*), making multiple dimensions interact simultaneously. Interactions between bacteria in the microbiome also have functional consequences (*3, 27–31*) and are prevalent in higher-dimensions (*3, 31*), where community assembly may introduce groups of species in different combinations e.g. in a fecal transplant.

Interactions in higher dimensions could change the topography of the fitness landscape (*31*), and their relative importance is unknown. To various extents, current approaches are limited in their ability to discern the topography of interaction landscapes in high dimensions due to (*i*) sign epistasis, which does not generalize well to more than two dimensions, (*ii*) a narrow ability to account for genomic context, and (*iii*) statistical considerations of the false discovery rate due to multiple testing (*32, 33*). Several different concepts of epistasis exist in the literature (*34*). However, standard epistasis frameworks often rely on parameter fitting, which brings along additional constraints (*35*). “Circuits”, which can describe all possible epistatic interactions (*33, 36*), introduce false discovery rate challenges.

Here we develop a new formulation of the *fitness cube* concept (*10, 13, 14*), where each biological entity (gene, organism, environmental factor) is a separate dimension (Fig. 1a). Because the biological entities are discrete (i.e., either a bacterium is there or it is not), our framework is discrete too. The cubes represent the landscape for interactions and can be composed of many dimensions as *n*-dimensional hypercubes. We then develop *epistatic filtrations* to locate the epistasis on this fitness landscape. Our approach solves problems of context and sign while reducing multiple testing concerns, all in a parameter-free form that is consistent across many dimensions (Fig. 1, Box 1).

**Figure 1:**
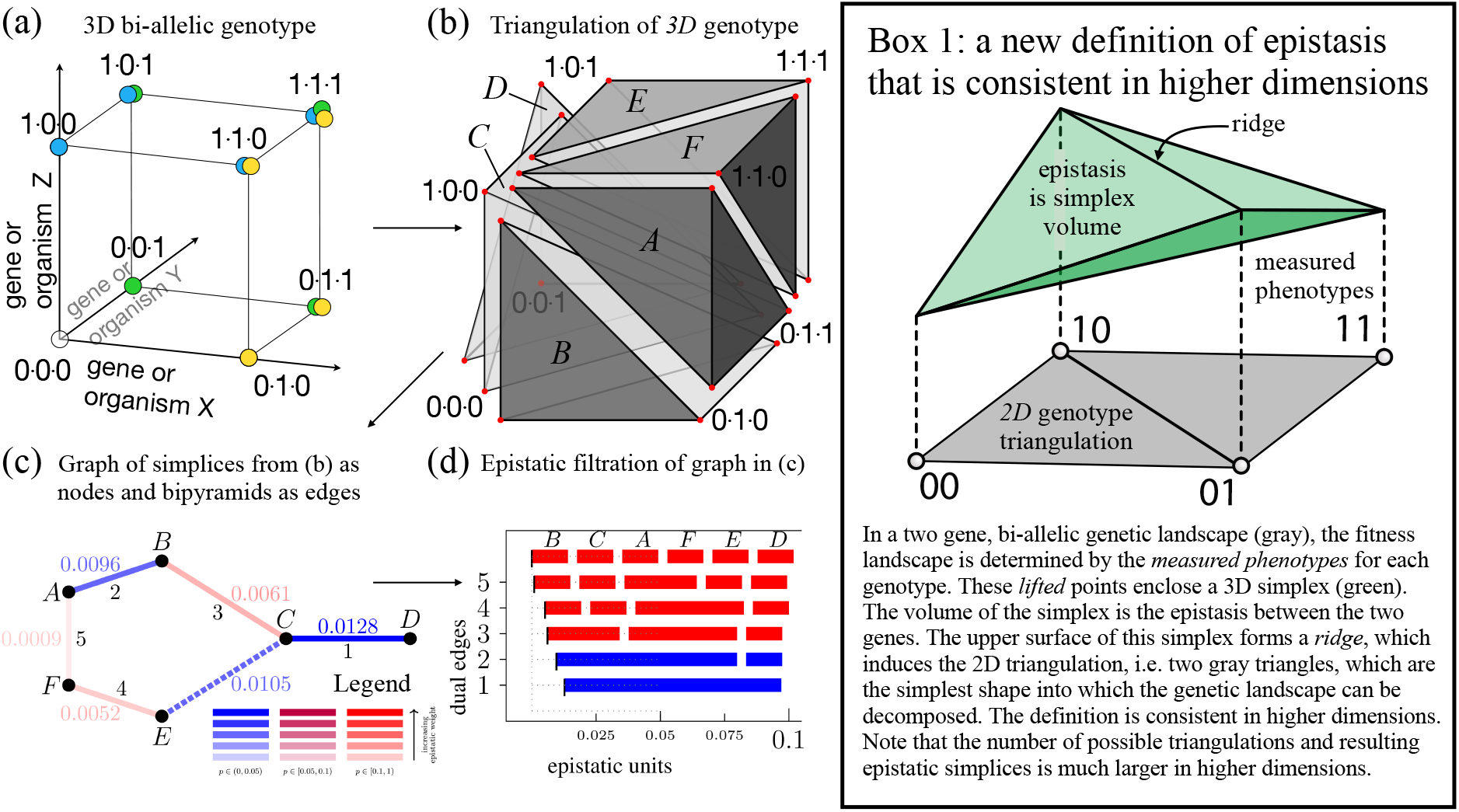
Filtrations describe epistatic topography. (a) Interacting biological entities, e.g. genes in a cell or bacterial strains in a microbiome, can be depicted as orthogonal dimensions in a unit cube, where vertices represent different genotypes or combinations of strains. (b) With 3 dimensions, the triangulation (*14*) of the fitness landscape produces *3D* simplices (labeled *A-F*) of the genotypes, and *4D* simplices of the fitness landscape (not shown) give the epistasis. (c) To map the global connectivity of the landscape, we merge adjacent simplices in a dual graph of the 3-cube triangulation, where nodes *A-F* are the simplices from (d) and the edges are the volumes of the bipyramids from the merges of neighboring simplices. The smallest bipyramid, edge 5, is formed first, followed by the next larger and so forth on up to the largest bipyramid, edge 1. The data set is from *Esherichia coli* mutations in *topA*, *spoT*, and *pykF* from (*37*). (c legend) Each dual edge has two parameters: its epistatic weight (indicated by shade) and its *p*-value (indicated by color). Black indices in (c) label the *critical* dual edges of 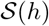, where critical indicates that loss of the edge leaves nodes unconnected to the graph. (d) The sequence of merges between adjacent simplices (reading from top to bottom) shown in the dual graph is depicted by the epistatic filtration. Epistasis of the merged simplex is indicated by the thin, black vertical hatch mark on the far left bar of each row. Total width of the bars is fixed. Note the non-critical C+E merge is not depicted in the filtration because those simplices are already merged with B, A, and F.

## 2 Results

### 2.1 Defining the shapes of fitness landscapes

We first decompose the fitness cube into its most elementary parts through a triangulation (Box 1, Fig. 1b). Triangulations are used e.g. in computer vision to decompose a surface, such as a human face, into discrete parts, which are triangles. Generalizing to higher dimensions, the triangles connecting genotypes are simplices (Fig. 1b). The volume of each simplex connotes the local steepness of the landscape (Box 1). To establish the global topography of the landscape we merge adjacent simplices in a stepwise manner such that flattest parts of the landscape are merged first and the steepest parts last (Fig. 1c). Each adjacent pair of merged simplices, *s* and *t*, forms a bipyramid, (*s, t*) through their shared face. The **epistatic weight** of (*s, t*) is

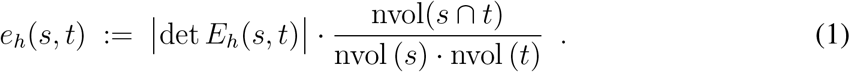

where *E*_*h*_(*s, t*) is the matrix specifying the vertices with their corresponding fitness phenotypes and nvol denotes the dimensionally normalized volume of the genotypes (Box 1; Appendix B1-B6). We use the notation

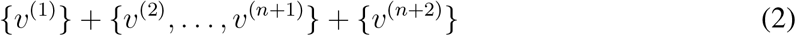

for the bipyramid (*s, t*), where the first and last vertices are the apices and the middle set forms the shared face. The *n*+2 genotypes of the bipyramid form a non-linear interaction of dimension *n* when *e*_*h*_(*s, t*) > 0.

We visualize the topography of the **epistatic landscape** by forming a **dual graph** of 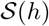, where the nodes are the maximal simplices and adjacent simplices form the dual edges. Blue edges indicate epistasis (Fig. 1c). The **epistatic filtration** of *h* (Fig. 1d) depicts the path from lowest to highest epistasis by merging adjacent simplices to form a connected **cluster** c.f. (*38*). In this sense, epistatic filtrations encode a global notion of epistasis in higher dimensions by connecting adjacent bipyramids. This method has many advantages over parameter fitting, including that it does not depend on the statistical constraints of determining a best fit.

Filtrations are also not constrained by the sign of epistasis, which depends on which genotype is considered *wildtype*, a somewhat arbitrary decision given varied ancestries (see Appendix B1). Studying adjacent simplices and their neighboring relationships, as we propose below, allows reconstruction of the fitness landscape and its epistatic properties in high dimensions. This process rests on the mathematical theory of linear optimization, convex polyhedra, and regular subdivisions (*38*).

We note that bipyramids account for the majority of genomic contexts (*38*), c.f. Table S1. Furthermore, the location(s) of inferred epistasis is robust to the choice of triangulation 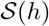 (*38*).

### 2.2 An evolutionary genetics example of epistatic filtrations

To illustrate our approach, we examined an existing data set from Lenski’s (*39*) classic experimental evolution of *Esherichia coli*, in a set of strains with each combination of five beneficial mutations (*37*) (Fig. 2a). We first examine *n* = 3 loci, corresponding to biallelic mutations in *topA*, *spoT*, and *pykF* (Fig. 1c,d). Epistasis was generally low in magnitude (*37, 40*), and occurs in two ways: (*i*) either from merging groups of groups of simplices (c.f. BC + AFE in line #2 of Fig. 1d), which indicates a complex interaction, or (*ii*) from merging a single simplex, c.f. D, with the aggregated rest of the simplices (c.f. line #1 of Fig. 1d), much like a dominant effect in the NK model (*14*). This second way is consistent with a fitness landscape distortion, which occurs when certain mutations influence the interactions of many other genes (*41*). Geometrically, such a distortion constitutes a vertex split (*42*). We next add a fourth biallelic mutation, in the *glmUS* locus (Fig. 2b,c), encoding peptidoglycan availability, which is an essential component of the cell wall.

**Figure 2:**
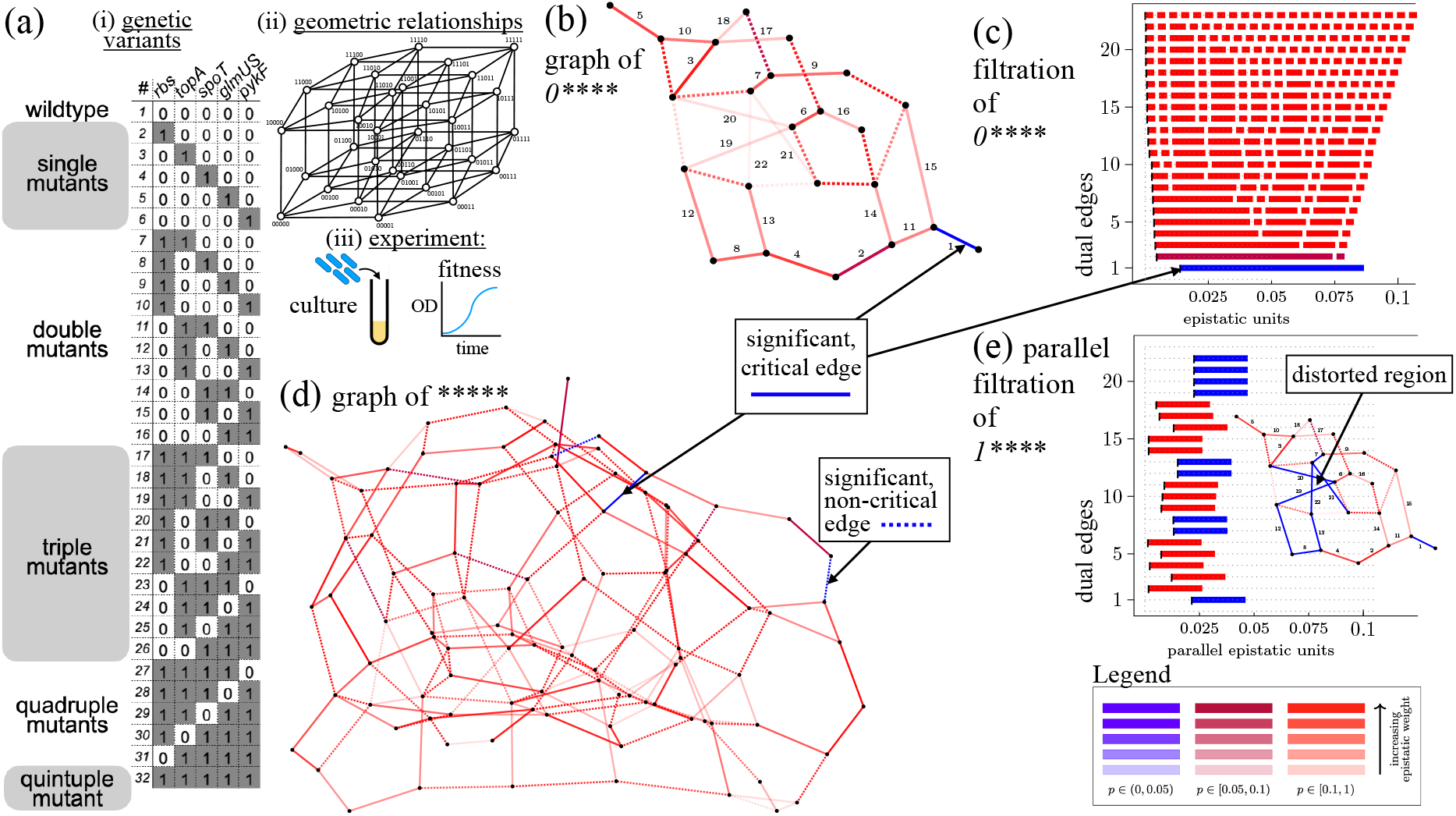
*E. coli* evolution is guided by epistatic landscape distortions. (a) (*i*) *E. coli* mutants examined (*37*), (*ii*) their geometric relationships, and (*iii*) experimental approach to measure fitness. (b) Edge labeled dual graph and (c) epistatic filtration restricted to *n* = 4 mutations in *topA* (locus 2), *spoT* (locus 3), *glmUS* (locus 4) and *pykF* (locus 5). Locus 1, *rbs*, is fixed 0 (*wildtype*). Note that the left edge of the bars in (c) indicates there is very little epistatic weight added to the filtration except for the final merge, where the single genotype 00001 gives weight to the entire filtration. This final interaction corresponds to the vertices {00001} + {00000, 01001, 00101, 00011} + {00010}. (d) Dual graph for the complete Khan data set. Black indices in (b) label the critical dual edges of 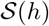. (e) In the parallel filtration, for 1 ****, where the *rbs* mutation is present, the landscape is disorted by a concentrated area of higher epistasis. Inset: graph in (b) recolored with weights from (e).

The filtration reveals a smooth, additive landscape with one dominant cell where epistasis arises only in the final merge of the filtration (Fig. 2c), meaning the epistatic topography of the entire landscape (Fig. 2d) rests upon the single vertex, 00001, *pykF*. While the previous analysis detected a significant, marginal effect of *pykF* (*37*), filtrations reveal the geometric structure in terms of which specific combinations of loci are responsible for the effect (Fig. 2e): the interaction between the *glmUS*, {00001}, and *pykF*, {00010}, genes requires the context of four loci, {00000, 01001, 00101, 00011}, yet it involves only up to double mutants, suggesting high dimensional epistasis that arises from lower dimensional interactions (Fig. 2c). This conclusion is consistent with recent genome-wide work on trans-gene interactions (*19*), suggesting that complex traits may arise from genome-wide epistasis, where each mutation’s contribution to the trait depends on the context of other mutations.

We introduced **parallel transport** (*38, §6.6*) to give a geometric measure of context-dependence for the same set of loci with different bystanders (e.g. species or genes) (see Fig. S1), previously examined by conditional or marginal epistasis (*43*). Examining the Khan data with and without the *pykF* mutation (*37*) (Fig. S2) showed increased significance in 9 out of 20 of the dual edges (Fig. S2), when *pykF* was mutated. Examining the restoration of *pykF* (Fig. S3), only 3 of 22 edges changed significance and just one critical edge lost significance, indicating that the epistasis in this case occurs because the mutation causes new interactions. Thus, the *pykF* mutation appears to enable further evolution during the Lenski experiment (*39*) by distorting the epistatic landscape. *rbs* also generates distortions (Fig. 2e), which can be visualized as a concentrated region of epistasis on the dual graph (Fig. 2e Inset). We found similar features in another genetic data set for the *β*-lactamase enzyme (*44*) (Appendix B7). Filtrations can thus reveal the specific geometric structure of both the interactions and the context they rely upon.

### 2.3 Lactobacilli produce microbiome distortions

Up to this point, we have focused on genetic epistasis, but our framework is generalizable to interactions of environmental parameters, including the gut microbiome, for which a framework to identify complex interactions is greatly needed. Like the genome, which is composed of many genes that interact to determine organismal fitness, the microbiome is also composed of many smaller units (bacterial species in this case) that affect host fitness. Hosts are known to select and maintain a certain core set of microbes (*45, 46*); the interactions of these bacteria can affect host fitness (*3*); and it is debated to what extent these interactions are of higher-order, c.f. (*28*). While vertebrates have a gut taxonomic diversity of ≈ 1000 species, precluding study of all possible combinations, the laboratory fruit fly, *Drosophila melanogaster*, has naturally low diversity of ≈ 5 stably associated species (*47*).

We made gnotobiotic flies inoculated with each combination of a set of *n* = 5 bacteria (2^5^ = 32 combinations) that were isolated from a single wild-caught *D. melanogaster*, consisting of two members of the *Lactobacillus* genus (*L. plantarum* and *L. brevis*) and three members of the *Acetobacter* genus (Fig. 3a). We measured fly lifespan, which we previously identified as a reproducible phenotype that is changed by the microbiome (*3*). Overall a reduction of microbial diversity (number of species) led to an increase in fly lifespan as with a taxonomically similar set of bacteria we examined previously, which came from multiple hosts (*3*).

**Figure 3:**
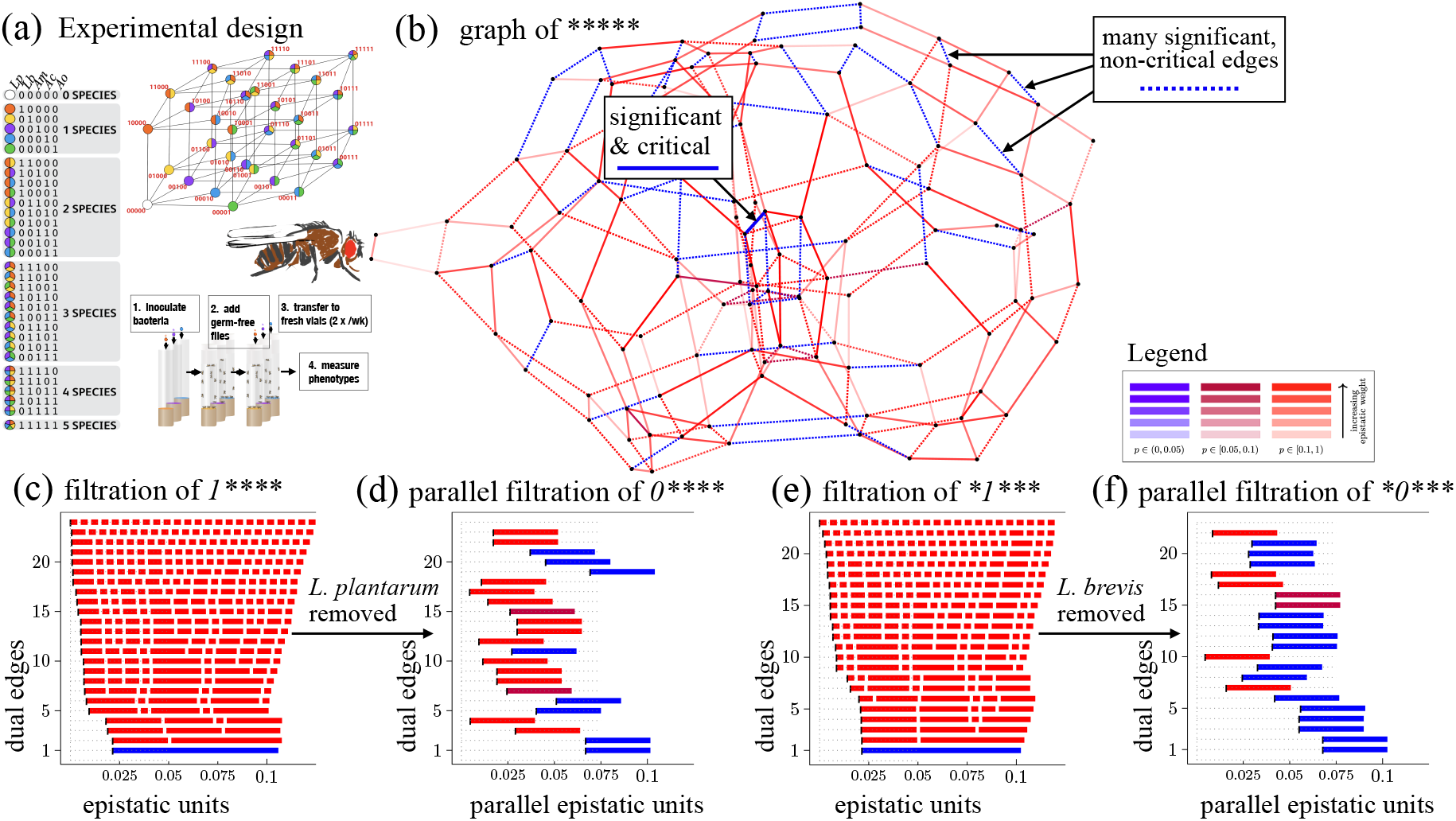
Loss of lactobacilli causes global distortion of the microbiome epistastic landscape. (a) Experimental design for Eble and Gould (*3*) microbiome manipulations in flies. (b) Full graph of for ***** the Eble data. (c) Filtration of 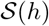 for the 4-face, 1****, of Eble data, where *L. plantarum* is present, indicates epistasis where two clusters of maximal cells merge. (d) Parallel filtration with *L. plantarum* removed shows a landscape distortion. (e) Filtration for *1***, where *L. brevis* is present has similar structure to 1****. (f) Parallel filtration with *L. brevis* removed shows a landscape distortion.

Epistasis was concentrated at the center of the dual graph (Fig. 3b,c), with significant, non-critical edges distributed throughout the graph (Fig. 3c). Examining the parallel transport, we found that the *Lactobacilli* drive changes in the global structure (Fig 3d,e). In 46 out of 128 (36%) interactions, significance changed due to adding or removing a *Lactobacillus* (Fig 3c-f, S7, S8). These changes in significance primarily derive from non-significant interactions when *L. brevis* is present that become significant when it is removed and vice versa, indicating *L. brevis* suppresses epistatic interactions that affect fly lifespan.

Microbiome abundances could drive the effects on host lifespan, however, comparing the epistatic landscapes for CFUs and lifespan, we found that only 2 of 99 dual edges were significant for both the bacterial abundance and fly lifespan data sets (Fig. S9, S10, S11, S12, Tables S2, S3, S4, S5), and there was a lack of correlation between the epistatic weights of the bipyramids (Spearman rank correlations: *p* = 0.7, *p* = 0.5, *p* = 0.3, and *p* = 0.3 respectively). This discord between the epistatic landscapes for microbiome fitness and host fitness could e.g. diminish the rate of co-evolution.

### 2.4 Interactions are sparse in higher-dimensions

We used epistatic filtrations to systematically evaluate the prevalence of higher-order interactions as a function of the number of dimensions. Critical, significant, higher-order interactions were less frequent than pairwise interactions (*p <* 10^−6^, *Z-test*) for each of the Khan, Eble, and Gould data sets, with a decreasing probability as a function of the face dimension (Table 1). This occurs for three primary reasons. First, the degrees of freedom increase in higher dimensions. Second, the probability of selecting a significant interaction from the set of all possible interactions decreases because the total number of interactions increases with increasing dimensions. Finally, the absolute number of significant interactions decreases in higher dimensions (Table 1), meaning they are biologically less prevalent. Overall, ≈ 10% of possible dual edges were significant at higher order, with ≈ 1% significant for *n* = 5 dimensions (Table 1), suggesting limits to the dimensions of biological complexity.

**Table 1:** Prevalence of interactions at different levels of complexity in genetics and microbiome data sets. Significant versus all critical dual edges (*p <* 0.05).

| Interaction dimension | Dataset:<br>Khan | Dataset:<br>Eble | Dataset:<br>Gould |
| --- | --- | --- | --- |
| 2: | 20/80 (25%) | 24/80 (30%) | 22/80 (28%) |
| all higher order: | 29/508 (5.7%) | 58/540 (10%) | 21/520 (4.0%) |
| 3: | 21/194 (11%) | 35/199 (17%) | 14/194 (7.2%) |
| 4: | 7/214 (3.2%) | 22/226 (10%) | 6/216 (2.7%) |
| 5: | 1/100 (1.0%) | 1/115 (0.8%) | 1/110 (0.9%) |
| total: | 49/588 (8.3%) | 82/620 (13%) | 43/600 (7.1%) |

We note that these fewer interactions in high dimensions can and do impact fitness. For example, the two top 4-dimensional interactions in the Eble microbiome data produce a combined 9% effect on fitness (see edges 1 and 2 in (Fig. 3)) with the largest maximal cell accounting for ≈ 5%. The relative sparsity makes for a tractable number of these interactions, where we may eventually determine the mechanisms, and filtrations provide a way to identify these.

### 2.5 Higher-order interactions can arise from lower-order interactions

Non-linearities of lower-order interactions can produce interactions in higher dimensions (*40*). In examining the higher-order epistasis present in our data sets, we noted that the clusters where significant epistatic weights occur are often preceded by clusters with nearly significant epistatic weights in lower dimensions (Fig. S4). We developed a graphical approach to distinguish these interactions from those that arise *de novo* (Fig. S20b,c; Appendix B11).

Several higher-order interactions in the Gould and Khan data could not be attributed to lower-order effects (Table S6). In particular, they could not be detected from pairwise interactions between loci, (c.f. Fig. S20c). As we noted, the 4-dimensional interaction in the *E. coli* evolution experiment involved loci with two genes (Fig. 2), whereas in the microbiome, interactions involved loci with four species, indicating different underlying geometries at these different scales of biology (Table S6).

## 3 Discussion and Conclusions

From an evolutionary perspective, the Red Queen’s hypothesis emphasizes how conflicts with other organisms can drive continuous genetic innovation (*48*). We find that epistasis in higher dimensions generates fitness landscape distortions, which could continuously change the fitness landscape to fuel new genomic innovation even in a static environment. This could partially explain the observation of continuous diversification in long term evolution experiments (*49*). In higher dimensions, we lack simple terminology to describe the many types of interactions that may occur, whether between quadruples and singles, pairs and triples, or different genetic backgrounds. We found that biologically-significant interactions in four and five dimensions are sparse and often rooted in lower order, meaning that a limited number of such interactions exist. This extends to higher dimensions the trend that 3-way interactions are often predicted from 2-way interactions (*2, 3, 28*). However, our finding that key genes and species cause distortions emphasizes the need to identify the significant higher-order interactions from the vast number of possible ones, a task that epistatic filtrations enable.

This geometric approach could be extended, e.g. to GWAS (*15, 19, 50*), ecosystems (*4, 5*), or neuronal networks (*51*), to discover non-additive higher-order structures at different scales. It should be noted that the polyhedral geometry methods for analyzing epistasis deserve to be developed further from the mathematical point of view. We believe that concepts of curvature for piecewise linear manifolds will be useful (*52*).

## 4 Acknowledgements

The authors acknowledge L.J. Holt and O. Brandman for insightful comments on the manuscript. Research by M.J. is carried out in the framework of Matheon supported by Einstein Foundation Berlin. Further partial support by Deutsche Forschungsgemeinschaft (SFB-TRR 109: “Discretization in Geometry and Dynamics” and SFB-TRR 195: “Symbolic Tools in Mathematics and their Application”. W.B.L. acknowledges NIH grant DP5OD017851, NSF IOS award 2032985, and the Carnegie Institution for Science Endowment.

## 5 Competing interests

The authors declare no competing interests.

## 6 Supplementary Materials

Materials and Methods

Fig S1 – S22

Tables S1 – S9

## A Materials and Methods

### A.1 Fly husbandry

Flies were reared germ-free and inoculated with one combination of bacteria on day 5 after eclosion. *N* ≥100 flies were assayed for lifespan in *n*≥5 independent vials per bacterial combination for a total of 3200 individual flies. Food was 10% autoclaved fresh yeast, 5% filter-sterilized glucose, 1.2% agar, and 0.42% propionic acid, pH 4.5. Complete methods are described in Gould *et al* (*3*).

### A.2 Bacterial cultures

Bacteria were cultured on MRS or MYPL, washed in PBS, standardized to a density of 10^7^ CFU/mL and 50 μL was inoculated onto the fly food. Strains are indicated in Table S7. See Gould *et al* (*3*) for complete methods.

### A.3 Genetics data

Existing genetics data sets were gotten from Sailer and Harms 2017 (*40*) github repository (https://github.com/harmslab/epistasis) or from Tan *et al* (*44*).

For the Khan data in Fig. 2, the fitness function *h* is defined for (b) by assigning the following normalized values to the 16 genotypes:

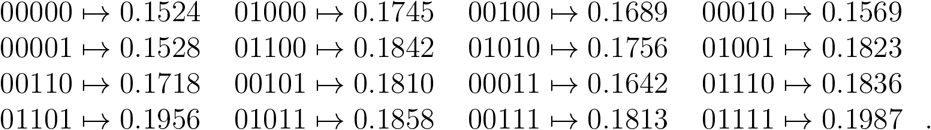

The Tan data set is different from the other fitness values in that only median and mean values are given, meaning we cannot compute *p*-values to assess the statistical significance. The fitness values are minimum inhibitory concentrations of antibiotics from a well-standardized assay with little experimental variation. Thus, the measurements and our analysis are believed to be robust. We note that the regular subdivision resulting from the corresponding height function of [0, 1]^5^ is degenerate in the sense that it is not a triangulation. This degeneracy arises because the data are discrete antibiotic concentrations with 24 possible values. The repetition of exact values in several cases means a triangulation does not occur. We extended our methods to this degenerate case by restricting the analysis to the faces that do have a triangulation, broadening the application of our approach. We focused on the piperacillin with clavanulate data from (*44*) as it is the better behaved.

### A.4 Computational analysis

The filtrations code is available as a polymake (*53*) package (cf. https://github.com/holgereble/EpistaticFiltration) and the analysis pipeline is available as a `jupyter` notebook.

## B Terminology

**Loci** (singular **locus**) refer to individual sites in the genome where a mutation may occur, or in the microbiome sense, a locus is a particular bacterial species. We write [*n*] := {1, . . ., *n*} for the set of all loci.

**Genotypes**, *v* = (*v*_1_, . . ., *v*_*n*_), are vectors of loci with 0/1-coordinates that form points in some fixed Euclidean space ℝ^*n*^, where *n* is the number of genetic loci or bacterial species considered. In this article we focus on **biallelic** *n*-locus systems, i.e. genotype sets of the form *V* = {0, 1}^*n*^ where *n* is the number of loci and each locus is either 0, absent, or 1, present. For instance, *v* = (1, 0, 1) denotes a genotype in a 3-locus system ℝ^3^, where the first and third loci are mutant and the second is wild type. The set of all genotypes will be denoted by *V*. The convex hull *P* := conv(*V*) of all genotypes is called the **genotope**. In our setting *P* is the *n*-dimensional unit cube [0, 1]^*n*^ (c.f. (Fig. S21) for a 2*D* projection of [0, 1]^5^).

A **fitness function** (also called **height function**) associates to each genotype *v* ∈ *V* a quantified **phenotype** describing the impact of the genotype on the organism. For example, if the measured phenotype is fitness, *h* encodes the reproductive output of the genotype.

The **fitness landscape** is the pair (*V, h*), which defines the fitness *h*(*v*) for each genotype *v* ∈ *V*. Let *v* = (*v*_1_, . . ., *v*_*n*_) ∈ *V* be a genotype. Then its **lift** is given by (*v, h*(*v*)) = (*v*_1_, . . ., *v*_*n*_, *h*(*v*)) ∈ ℝ^*n*+1^.

A set of points *W* = {*w*^(1)^, . . ., *w*^(ℓ)^} is **affinely independent** if for all real scalars *λ*_*i*_ satisfying 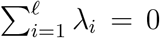 the condition 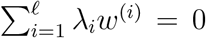 forces *λ*_*i*_ = 0 for all *i* ∈ {1, . . ., *ℓ*}. Otherwise *W* is **affinely dependent**.

An **interaction** with respect to a fitness function *h* occurs between a collection of *k* + 2 affinely dependent genotypes *v*^(1)^, . . ., *v*^(*k*+2)^ ∈ *V* ⊂ ℝ^*n*^, for *k* ≤ *n*, whose lifts are affinely independent points in ℝ^*n*+1^. This is in line with the standard concept of additive epistasis. The number *k* is the **dimension** of the interaction; throughout we assume that *k* ≥ 2.

Let *U* = {*v*^(1)^, . . ., *v*^(ℓ)^} be a set of genotypes. Its **support** is the set

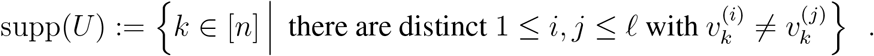

That is, the support is the set of loci where at least two of the given genotypes differ. For example, if *n* = 3 and *U* = {(0, 0, 0), (1, 0, 1), (1, 0, 0)} then supp(*U*) = {1, 3}.

The number of loci that vary (0 vs 1) in the support is called the **order** of an interaction; this definition agrees with, c.f., (*54*): “We designate interactions among any subset of *k* mutations as *k*th-order epistasis.”. We give two examples: First, let *n* = 2 and *U* = {(0, 0), (0, 1), (1, 0), (1, 1)} = *V* such that *U* is an interaction with respect to some fitness function. Then *U* is an interaction of dimension 2 and order 2. Second, let *n* = 3 and *U* = {(0, 0, 0), (0, 1, 1), (1, 0, 0), (1, 1, 1)} such that, again, *U* is an interaction with respect to some height function. Then the dimension is 2 and the order is 3. In general, the order is at least as large as the dimension, but the two quantities may differ. We say that genes (corresponding to loci) **interact** if they form the support set of an interaction of genotypes.

**Remark.** The dimension *k* of an interaction *v*^(1)^, . . ., *v*^(*k*+2)^ with respect to some fitness function agrees with the dimension of the affine span of the given points in ℝ^*n*^. This can be seen as follows. By definition the lifted points (*v*^(1)^, *h*(*v*^(1)^)), *. . .,* (*v*^(*k*+2)^, *h*(*v*^(*k*+2)^)) are affinely independent in ℝ^*n*+1^. So their affine span has dimension *k* + 1. As *v*^(1)^, . . ., *v*^(*k*+2)^ are affinely dependent, the dimension of their affine span is at most *k*. Now the affine dimension can only increase by at most one if one coordinate is appended.

### B.1 A primer on epistatic filtrations

We first explain the biallelic case with *n* ≥ 2 loci. In the geometric framework (*33*), two interacting loci give rise to four possible genotypes, which form the vertices of a square and may be written as vectors of zeros and ones, indicating the absence (0, wildtype) or the presence (1, mutant) of each locus respectively (Fig. 1b) (*33,38*). The measured phenotypes lift the genotype vertices into 3-space, and there is epistasis corresponding to the volume of the simplex enclosed by the lifted points (see blue simplex in Fig. 1b). Geometrically, the four genotypes involved are fully symmetric, meaning that the sign of the epistasis for *n* = 2 is relative to the choice of a coordinate system. Thus, the sign of epistasis depends on which genotype is considered wild-type. By considering the simplex volume rather than the fold of the upper shell of the simplex, epistatic filtrations do not specify a sign and thus avoid this caveat. However, directionality is considered by parallel transport (see later section). Returning to our explanation, by taking the upper convex hull of all 2^*n*^ lifted points and projecting back onto the genotope [0, 1]^*n*^ we induce a **subdivision** 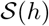; cf. (*38,55, §2.1*), into **maximal cells** (Fig. 1b). Generically, every maximal cell of 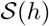 is an *n*-dimensional simplex, which is the convex hull of (*n* + 1) affinely independent genotypes (Fig. 1c). Importantly, these *n*-dimensional simplices are the most elementary parts into which a fitness landscape can naturally be decomposed.

Our framework generalizes to higher dimensions through a geometric shape called a **bipyramid**, where two satellite vertices, each the apex of one pyramid, are joined to a common set of base vertices. The satellites correspond in the 2*D* example (Fig. 1b) to 00 and 11 and the base to 10 and 01. This is naturally associated with 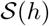, set up by the **ridge** (Fig. 1b). For an ordered sequence of *n* + 2 genotypes (*v*^(1)^, *v*^(2)^, . . ., *v*^(*n*+2)^) we let

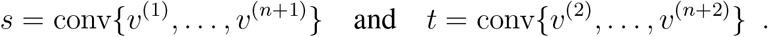

In other words, *s* and *t* form convex hulls. We call such a pair (*s, t*) a bipyramid with vertices *v*^(1)^, *v*^(2)^, . . ., *v*^(*n*+2)^. Then we can find the volume of the lifted bipyramid by forming the (*n* + 2)×(*n* + 2)-matrix

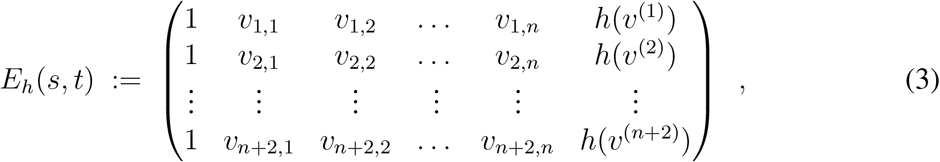

where *v*_*i,*1_, *v*_*i,*2_, . . ., *v*_*i,n*_ are the coordinates of *v*^(*i*)^ ∈ ℝ^*n*^. The **epistatic weight** of the bipyramid (*s, t*) is

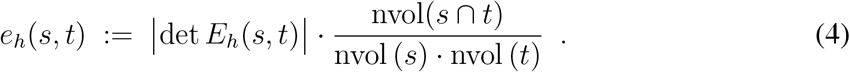

Here nvol denotes the dimensionally normalized volume. The quantity nvol(*s*∩*t*) is the relative (*n*−1)-dimensional normalized volume of the **ridge** of the bipyramid, given by the intersection *s* ∩ *t* = conv(*v*^(2)^, . . ., *v*^(*n*+1)^). We use the notation

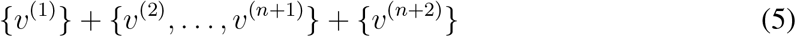

for the bipyramid (*s, t*), where the first and last vertices are the satellites and the middle set forms the base. Now the *n* + 2 genotypes of the bipyramid form an interaction of dimension *n* when *e*_*h*_(*s, t*) > 0.

In our regular triangulation 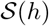, the two *n*-dimensional simplices, *s* and *t*, are **adjacent** because their intersection *s* ∩ *t* is a common face of dimension *n* − 1.

### B.2 Constructing a filtration from the epistasis of adjacent simplices

We visualize the topography of the **epistatic landscape** by forming a **dual graph** of 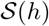, where the nodes are the maximal simplices and adjacent simplices form the dual edges. A rugged path is one with more blue edges (Fig. 1d). To each such dual edge we associate an epistatic weight and a label (Fig. 1c, epistatic weights are in shades of blue and red, while labels are in black). In this way, we construct an epistatic landscape that corresponds to the underlying fitness landscape with the ruggedness specified along the dual graph. The **epistatic filtration** of *h* (Fig. 1e) depicts the path from weakest to highest epistasis by merging adjacent simplices. These diagrams summarize the information contained in epistatic weights and dual graphs, and facilitate comparisons across data sets. But there is important new information contained in epistatic filtrations, which is not directly visible from the dual graph and its epistatic weights. Indeed, a step in the epistatic filtration merges adjacent simplices. We build the complete fitness landscape by stepwise merging of maximal cells, starting from the lowest epistatic weight and stepwise merging adjacent simplices to form a connected **cluster** c.f. (*38*). In this sense, epistatic filtrations encode a global notion of epistasis in higher dimensions by connecting adjacent bipyramids.

To see this, notice that each row of the diagram has a number of bars and a black leftmost line. In the top row the black line marks the epistatic weight of zero (*x*-coordinate). Each bar is red and corresponds to one maximal simplex of 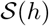. In the second row (counting from the top), we see three things: (1) the value of the lowest epistatic weight moves the *x*-coordinate of the black line slightly to the right. (2) The two maximal simplices of 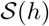 corresponding to this epistatic weight are merged into one. These correspond to the two bars in the previous row above the new, longer bar in the row. The lengths of the other bars remain unchanged but are shifted horizontally by the epistatic weight in (1). (3) The statistical significance of the epistatic weight giving rise to the merging step, encoded by the colors of the bars; cf. Section B.4.

The merging procedure is then repeated for each pair of maximal simplices arising in each epistatic weight until one reaches the highest epistatic weight and the last maximal simplex of 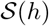 to be merged with the rest. In this way the indentation of the bar charts increases from top to bottom. The total width of the bars stays constant throughout.

Importantly, in the epistatic filtration diagram, not every merging step is displayed; e.g., in Fig. 1d there are fewer rows than dual edges in Fig. 1c. This is because some steps do not change the resulting fitness landscape (no actual new portion is merged to the previous one). The reported steps are only the ones increasing the connected components of the fitness landscape obtained from the previous merging steps. The epistatic weights corresponding to these steps are the edges in the dual graph which we call **critical** in (*38, §.3.2*).

### B.3 Normalized epistatic weights

To gain a perspective on the generality of higher-order interactions, it is desirable to compare epistatic landscapes. Different phenotypes have different metrics, making comparisons difficult for current approaches to epistasis. Filtrations are well-suited in this sense. Scaling the height function *h* by a positive constant does not change the regular triangulation, and thus it does not change the dual graph. In order to compare different data sets, we scale the height function to Euclidean norm one. The epistatic weights are scaled accordingly. The resulting **normalized epistatic weights** are measured in **epistatic units**, giving a generalized metric for epistasis.

Measuring the effect of context on epistatic interactions is also desirable, e.g. to detect the marginal or conditional effects of a locus (*37*), and these are a natural feature of filtrations. If we fix some *k* loci and let the remaining *n* − *k* loci vary, we obtain a height function, which is **restricted** to a face of the genotope [0, 1]^*n*^. That face has 2^*n*−*k*^ vertices, and it is an isomorphic copy of the cube [0, 1]^*n*−*k*^. For instance, if *n* = 5 and we fix the first and the fourth locus to 0, we obtain a 3-dimensional face, which we denote 0**0*. That is, such a face is written as a string of *n* symbols in the alphabet {0, 1, *}, where 0 or 1 mark the fixed choices, and * stands for variation. The number of * symbols equals the dimension of the face. Triangulations, their dual graphs, epistatic weights, etc. are well-defined for height functions restricted to faces. This aspect of the theory allows the study of conditional epistatic effects.

### B.4 Statistics of epistatic weights

We developed a statistical test to quantify the significance of an interaction associated with a fixed bipyramid; cf. (*38, §4.2*). Here we assume that *h*(*v*) is the mean value of the individual phenotype measurements for some number of replicated experiments for the fixed genotype *v*. To each dual edge we associate a *p*-value, which is independent of the epistatic weight normalization. If that *p*-value is below 0.05 we call that dual edge **significant**. It is useful to also consider *p*-values, which are slightly higher because one can use the shape of the landscape to identify interesting locations for further statistical analysis. To this end we call a dual edge **semi-significant** if 0.05 ≤ *p <* 0.1.

While it may be possible that this approach misses some biologically relevant interactions (e.g. if they do not correspond to a bipyramid selected by our method), those interactions that we identify carry information that is robust and supported by a statistical model. The fact that not all possible interactions can be approached is an inevitable consequence of the higher dimensional nature of fitness landscapes, also reflected by a very high number of possible regular triangulations of [0, 1]^*n*^. That number equals 74 for *n* = 3 and 87,959,448 for *n* = 4, whereas the precise numbers for *n* ≥ 5 are unknown; cf. (*55, §6.3*). Thus, filtrations use the data to greatly condense the number of possible interactions considered.

The bar colorings in the filtrations of epistatic weights, as in (Fig. S4), reflect the outcome of multiple simultaneous statistical tests (one for each epistatic weight) (*38*).

Significant dual edges at *p <* 0.05 are shown in blue, 0.05 ≤ *p <* 0.1 in purple, and *p* ≥ 0.1 in red.

It may happen that a triangulation has a significant dual edge, which is not critical, whence it does not show in the epistatic filtration. In that case the next critical dual edge becomes blue; so a filtration encodes all significant interactions found by our method.

**Remark.** By funneling the analysis through the concept of regular triangulations our approach pre-selects interactions, which are most relevant with respect to fitness (*38, §2.2*). Via this major deviation from (*33*) we are able to detect interactions in many data sets, which are biologically plausible; this suggests strongly that our method is particularly good at avoiding false positives. Future work will investigate the relationship to other methods from statistics and signal processing. While most of this is beyond the scope of the present study, in Appendix B12 we offer a first step by comparing with traditional linear regression approaches.

### B.5 A synthetic experiment examining how epistatic weights change as a function of the interaction order

Our method calculates significance of detected interactions and normalizes the epistatic weight to the volume of the unit cube of the same dimensionality. We used synthetic data to analyze the method performance. We first examined 468 synthetic filtrations over the 4-dimensional cube, producing 10011 critical dual edges. We found that the epistatic weight is indeed constant as a function of the interaction order, see (Fig. S19a). This indicates that the normalization method is effective. Furthermore, the number of significant interactions decreased as the standard deviation of the input data increased, indicating the statistical method is sensitive to noise, see (Fig. S19b).

### B.6 A microbiome example in dimension 4

Here *n* = 4, and the fitness function *h* is defined by assigning the following values to the 16 genotypes:

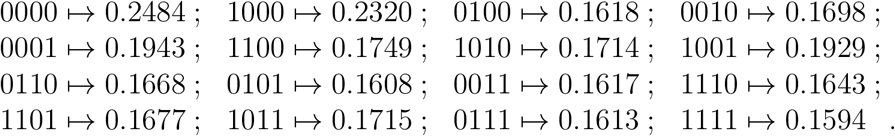

The vertices *U* := {*v*^(1)^, . . ., *v*^(6)^} ∈ *V* given by

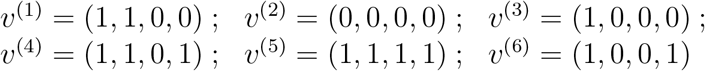

form a bipyramid (*s, t*) consisting of 4-dimensional simplices *s* and *t* as above. The simplices *s* and *t* correspond to nodes in the dual graph of 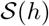 that share a dual edge recording their adjacency relation as indicated in (Fig. 3b).

In this situation, equation (4) reads

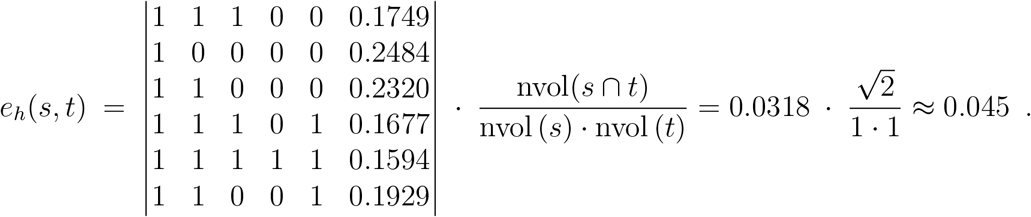

Since *e*_*h*_(*s, t*) > 0, the genotype set *U* defines a 4-dimensional interaction with full support {1, 2, 3, 4} and of order 4, according to our terminology of Section Terminology. With a *p*-value of 0.0005 < 0.05 the significance test established in (*38, §.4*) rejects the zero hypothesis for *e*_*h*_(*s, t*) and therefore proves the effect of the interaction *U* to be significant. We indicate this fact with the color blue both in the dual graph of 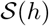 in (Fig. 3b) and in the epistatic filtration of *h* in (Fig. 3c).

This example illustrates the following fact of biological interest. For the bacterial combinations *v*^(1)^, *v*^(2)^, . . ., *v*^(6)^ fitness, given by the fitness function *h*, varies significantly in a non-linear way.

### B.7 The epistatic landscape within a single enzyme is rugged

As a point of comparison with the Khan data set, we re-analyzed data from a fully factorial 5-mutation data set in the *β*-lactamase gene, where each mutation is in a separate residue of the same enzyme (*44, 56*). Due to a lack of the raw replicate data, our computations are based on the reported mean values, and *p*-values are not calculated. The filtration holds a high magnitude of epistasis (Fig. S5, S6) compared with the Khan data set (Fig. S4, S2); note magnitude on the *x*-axis. The epistasis arises in many steps (note slope of filtration on left side; (Fig. S5, S6)), consistent with the low number of possible evolutionary paths observed by Weinreich (*56*), and distortions are apparent in the shifted magnitude of epistasis by parallel transport. Our geometric approach also reveals a tiered structure to the epistasis, c.f. the largest weight merges two clusters of simplices (Fig. S5, S6), indicating a more complex epistatic landscape than the Khan data set, where epistasis came from one individual simplex on the periphery of the dual graph.

Examining the filtration (Fig. 3d), the epistatic weight (i.e. magnitude) for the microbiome data generated ≈ 5% effect, roughly three times the weight in the Khan data and half that in the Tan *β*-lactamase landscapes (*44*) (c.f. *x*-axis between Fig. 3, S4, S5), indicating that the rugosity of microbiome interactions is comparable to genetic ones.

To further compare the global effect of context across different datasets, we developed a method to compute epistasis, based on the triangulation of dual landscapes, which we call the epistatic product [Appendix *Product model for epistasic landscape rugosity*] (Fig. S13, S14, S15, S16, S17, S18). The total epistasis was highest for the *β*-lactamase experiment (*44*), which carries much higher context-dependence than either the microbiome (*3*) or *E. coli* evolution data sets (*37*), indicative of overall high epistasis at the smallest, within enzyme, scale.

### B.8 Interactions are sparse in higher-dimensions

The prevalence and importance of higher-order interactions is debated, with some studies suggesting pairwise interactions predict the vast majority of interactions in complex communities (*28*), and others suggesting a large influence of context-dependent effects (*3*) (*57*), which would make higher-order interactions unpredictable. As we showed in the previous section, few such interactions are biologically meaningful in the context of fitness.

This limitation on epistasis in higher dimensions could arise due to e.g. limited phenotypic dimensions where interactions can be detected or to a lower dimensional manifold that absorbs the majority of the effects (*58*) (e.g. lifespan and fecundity are anti-correlated, making fitness robust to changes in one or the other). Regardless, our analysis shows that significant epistatic interactions are increasingly sparse as the number of dimensions for interaction increase, indicating there exist some limits to biological complexity.

We analyzed the few higher-order interactions in greater detail using a geometric approach. As we noted previously, the interactions in the Khan genetic data (Table 1) are based on a vertex split of the genotype 00001, meaning that the entire epistatic weight of the landscape is balanced by a single maximal cell (Fig. 2).

In contrast, the epistatic filtration of the Eble microbiome data in (Fig. 3) has a much richer texture. There are two significant bipyramids

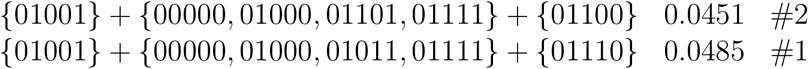

given with their epistatic weights and edge id’s, which form a cluster of interactions, indicating a larger topographic feature in the epistatic landscape that relates the interactions between *L. brevis* and increasing numbers of *Acetobacters*. Proximal to these significant cells are two cells with nearly significant statistical support:

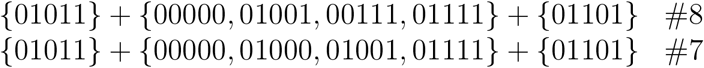

with their edge id’s (Fig. 3). This invites further research on the bacteria involved. For instance, the interactions could derive from metabolic crossfeeding between the *Acetobacters*, which produce many co-factors, and *L. brevis*, which produces lactate, stimulating *Acetobacter* growth (*59*). Note that the support sets of the bipyramids for all four interactions contain both the wild type 00000 and 01111, which are the maximum and minimum fitness respectively.

### B.9 Parallel transport of epistatic weights

The notion of parallel transport in a fitness landscape (*V, h*) was introduced in (*38, §6.6*) as a way to compare geometric and biological information between pairs of parallel facets of the convex polytope conv *V*. In this work, we extended that notion to include the case of two fitness landscapes, (*V, h*_1_) and (*V, h*_2_), associated to different generic and normalized height functions *h*_*i*_ : *V* → ℝ, *i* ∈ {1, 2}, defined on the same vertex set *V* = {0, 1}^*n*^ for some *n* ∈ ℕ. To enable meaningful comparisons, we assume that each *h*_*i*_ is normalized and that there is a larger fitness landscape (*W, h*) with a generic and normalized height function *h* : *W* → ℝ restricting to *h*_1_ and *h*_2_ on the parallel facets *V* in *W*, such that the partition of conv *W* induced by *h* is compatible with the one of conv *V* induced by *h*_1_, resp. by *h*_2_. In this setting, we define **normalized epistatic weights** as with Eq. (4) with *h* the normalized height function and *s, t* any adjacent simplices forming a bipyramid.

Parallel transports enable us to transport epistatic filtrations along the reflection map

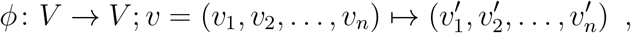

with *v′*_*i*_ = 1 − *v*_*k*_ if *i* = *k* and *v′*_*i*_ = *v*_*i*_ otherwise. More precisely, let 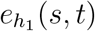 be the normalized epistatic weight associated to a bipyramid of 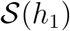 and let 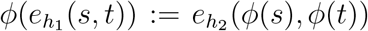 be the parallel normalized epistatic weight transported by *ϕ*. Then the filtration of normalized epistatic weights induces a filtration of parallel normalized epistatic weights. Additionally, to 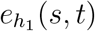 and to 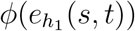 a *p*-value can unambiguously be associated (*38, §4.1-4.2*). Notice that by design epistatic filtrations for 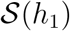 only show normalized epistatic weights associated to critical dual edges, defined as in (*38*). But normalized epistatic weights and their significance can be defined for all bipyramids including the ones associated to noncritical dual edges. This explains the labelling of the parallel transport tables below. There a row is numbered only if the bipyramid corresponds to a critical dual edge in the dual graph of 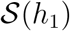. Noncritical dual edges whose normalized epistatic weight remains non-significant after the parallel transport are omitted. The normalized epistatic weight before (denoted by 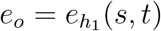) and after (denoted by 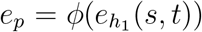) the parallel transport, as well as their *p*-values (denoted by *p*_*o*_ and *p*_*p*_) are also reported, as well as ratios of these quantities.

These parallel transport tables are linked to the epistatic filtration diagrams. Indeed, each numbered row in the table corresponds to the row in the epistatic filtration diagram with the black line set at *e*_*o*_. It also corresponds to the row with black line set at *e*_*p*_ in the parallel transported filtration diagram.

Recall from Section *Statistics of epistatic weights* that there may be dual edges of the triangulations which are significant but not critical. Since only the critical dual edges are labeled (by the row number in the epistatic filtration), in our tables for parallel transport these show up as unlabelled rows.

Examples for the parallel transport of epistatic filtrations are shown in Figures S1, S2, S3, S5, and S6. The magnitude of the epistasis in the left panels are roughly comparable between data sets due to normalization of the input data. Compare each left panel with its corresponding right panel to observe the relative change in epistasis in the parallel path. Larger changes in epistasis indicate stronger context-dependence of the interaction. For instance, in the first Weinreich comparison (Fig. S5), bar 10 in the right panel has a parallel epistasis greater than the original filtration on the left, indicating context-dependence.

### B.10 Product model for epistasic landscape rugosity

In this section we offer a new methodological framework to simultaneously study fitness landscapes associated to different height functions. We also provide a measure to quantify how much the height function of the combined fitness landscape differs from the sum of the height functions.

Let *U* and *V* be point configurations in ℝ^*m*^ and ℝ^*n*^, respectively. We think of these point configurations as two sets of genotypes, which may be distinct or not. If we have height functions *λ* : *U* → ℝ and *μ* : *V* → ℝ, then taking the sum *λ* + *μ* point-wise yields a lifting function of the product *U* × *V* ⊂ ℝ^*m*+*n*^. The cells of the regular subdivision 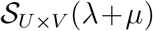 are products of cells of 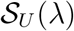 with cells of 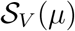. In particular, if *λ* and *μ* are generic, i.e., 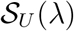 and 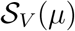 are triangulations, then the cells of 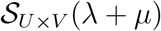 are products of simplices.

Now we consider an arbitrary height function *ν* : *U* × *V* → ℝ on the product of the point configurations. This yields height functions

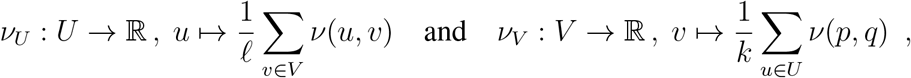

where *k* = #*U*, *ℓ* = #*V*, *u* is a vertex in *U* and *v* is a vertex in *V*.

Further we define

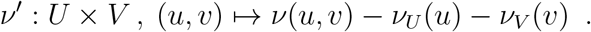

Observe that

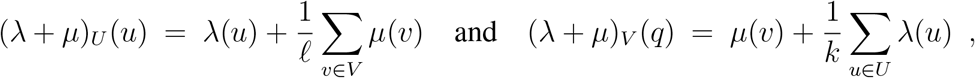

and (*λ* + *μ*)′ is the height function with constant value −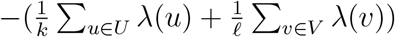. Thus *λ* + *μ* and (*λ* + *μ*)_*U*_ + (*λ* + *μ*)_*V*_ induce the same regular subdivision of *U* × *V*. Therefore, we propose to analyze the height function *ν′* to measure how much *ν* deviates from the sum of two height functions. We can use the techniques from our previous paper (*38*) and apply (all of) them to 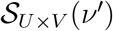 for any given *ν*. For instance, this allows to measure how independent two different height functions are on the same point set (this is the case *U* = *V*). We say that *ν* **decomposes as a product** if *ν′* = 0.

**Example 1.** If *U* = *V* = {0, 1} are the vertices of the unit interval then *U* × *V* are the vertices of the unit square [0, 1]^2^. Analyzing 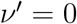 for any given height function *ν* on the four points (0, 0), (0, 1), (1, 0) and (1, 1) gives back the standard basic example of additive epistasis.

**Remark.** Two observations are in order: In (*38, §6.6*) we considered a version of parallel transport to compare epistatic effects, see also Appendix B9. The connection to the product model approach is as follows. Let *V* = {0, 1}^*n*^, i.e., the vertex set of the *n*-dimensional unit cube, be embedded twice, into a pair of parallel facets of the unit (*n*+1)-cube [0, 1] × [0, 1]^*n*^. This occurs in the product model with *U* = {0, 1}. If a height function *ν* on {0, 1} × *U* decomposes as a product then the parallel transport (in both directions) is trivial. Note that the number of dimensions is greater for the product model than for the parallel transport.

Additionally, observe that the product model differs from the marginal epistasis framework, which would produce a single number testing if the mutant changes one specific interaction between the genes.

#### B.10.1 Product model for the Khan data

To illustrate the product model consider the following example from the Khan data. We are interested in detecting if interactions between the *topA*, *spoT*, and *pykF* genes change when the *rbs* gene is mutated. To answer this question we let *U* and *V* be 3-cubes inside [0, 1]^5^ defined by three mutable loci, one for each of the above genes and indicated by *, and two fixed loci. The first fixed locus represents the *rbs* gene. It is not mutated in *U* and mutated in *V*. The height functions are compared over the three variable loci. Thus the filtration over the product model for *U* and *V* has four dimensions in this case. A computation reveals that there are no significant dual edges in the epistatic filtration on product model, see (Fig. S13). This indicates that the *rbs* mutant does not affect the interaction landscape.

### B.11 Meta-epistatic charts

This section deals with the question to which extent higher order epistatic effects are induced by lower dimensional ones or, put in other terms, which lower dimension epistatic effects can be seen in higher dimension. The **meta-epistatic chart** is a diagram drawn on top of the induced epistatic filtrations for some selection of faces of a fixed cube; higher-order interactions induced by lower order interactions are marked as corresponding.

In (Fig. S20b) and (Fig. S20c) we exhibit an example for the Eble data set, with 5 loci, where we take the five 4-dimensional faces 0****, *0***, **0**, ***0* and ****0 into consideration. Mathematically, these five 4-faces constitute the face figure of the wild type. Fix one 4-face, say 0****. The induced epistatic filtration on this face shows two blue bars corresponding to dual edges labeled 1 and 2. Each of them refers to the ridge of a bipyramid, which is a 3-dimensional simplex in this case. These two ridges may intersect certain 3-dimensional faces in the right dimension and thus may or may not descend to significant ridges within certain 3-dimensional filtrations. In case of an incidence with a lower dimensional significant ridge, the significant 4-dimensional effect is induced by a lower dimensional effect and one may picture this fact as a directed assignment pointing from the lower towards the higher dimensional interaction.

### B.12 Comparison with a simple linear regression approach

In the theory of fitness landscapes many linear regression approaches have been proposed to study higher-order interactions, c.f. (*21, 34, 40, 60*). In this section, we compare our epistatic weight method to an elementary regression approach using an example from the data.

The regression analysis we have in mind assumes that there is a linear relationship between the predictors *X*_1_, *X*_2_, . . ., *X*_*n*_ (one associated to each locus/dimension of the genotope) and response, or dependent, variables *Y* (associated to the biological measurements). That is, one assumes that *Y* = *f*(*X*_1_, *X*_2_, . . ., *X*_*n*_) + *ϵ* where *f* : ℝ^*n*^ → ℝ; (*X*_1_, *X*_2_, . . ., *X*_*n*_) → *β*_0_ + *β*_1_*X*_1_ + *β*_2_*X*_2_ + · · · + *β*_*n*_*X*_*n*_ and where *ϵ* is a random error term. The coefficients *β*_1_, *β*_2_, . . ., *β*_*n*_ are unknown but can be estimated by minimizing the sum of squared residuals associated to the observations pairs (*x, y*). These observations pairs consisting of a genotype and a measurement associated to it. Notice that more than one measurements are typically associated to a single genotype. With the coefficient estimates one can make predictions for the dependent variable via

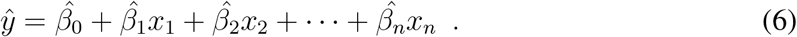

The hat symbol^indicates a prediction, for instance of *Y* on the basis of *x*_*i*_ = *X*_*i*_, or an estimate for an unknown coefficient.

Below, we are interested in the differences between the observed measurements *y* associated to the genotypes of [0, 1]^*n*^, expressed in terms of *x*_1_, *x*_2_, *. . . x*_*n*_ and the predicated values 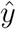 on the regression hyperplane (6). Notice that the regression analysis remains unchanged after normalizing the height function to Euclidean norm one. Additionally, computing residues for all replicated measurements (when provided) and then take averages builds on the assumption that measurements associated to different genotypes are statistically independent from each other. This assumption is consistent with the one underlying the computation of statistical significances for epistatic weights, following (*38, §. 4.2-4.3*).

**Remark.** In the regression setting of (6) there are hypothesis tests (like the *F*-statistic, *t*-statistics and *p*-value) to answer if at least one regression coefficient *β*_*j*_, 1 ≤ *j* ≤ *n* is nonzero, see for example (*61*). Such statistical approaches are different from the one in (*38, §. 4.2-4.3*), where other hypothesis tests for each epistatic weight were proposed.

#### B.12.1 Regression for Eble data

In the following, we perform a regression analysis focusing on the replicated measurements for the lifespan fitness landscape on [0, 1]^5^ obtained from Eble and subspaces thereof. Numerical measures of model fit (*F*-statistic: 2357, with *p*-value essentially zero, and for 3840 observations and 5 predictors) show that the multiple linear regression model can be considered to be appropriated for this data. Since the epistatic weights of the dual edges are close to zero (≤ 0.02) and are mostly not significant, the above regression analysis conclusion is in line with what we see from the filtration of epistatic weights associated to the same fitness landscapes, see (Fig. S22).

From this example we see that the regression approach provides some general information on higher-order interactions. However, without further assumptions, only one interaction formula is given in terms of a regression hyperplane (6) while the epistatic weight approach gives more fine grained information. This example also illustrate that when the regression model fits the data well (essentially the higher the *F*-statistics and the more coefficients in the hyperplane equation are significantly non-zero) the epistatic filtration has little horizontal shifts and few significant epistatic weights.

We now proceed repeating the above analysis on some of the bipyramids considered in the parallel analysis for the normalized lifespan Eble data. Regressing over bipyramid 23 in Table S8

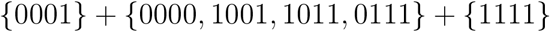

in 0**** and 1**** reveals that only two average residues over 0**** are non-zero (associated to the microbiomes 00000 and 00001), and only one is non-zero over 1**** (associated to the microbiome 10000). This confirms the two non significant epistatic weights over bipyramid 23 in Table S8.

**Remark.** If minimally dependent sets of points in the genotope are fixed, as in the epistatic weight approach, and one regresses above these points, then the corresponding regression hyperplanes equations are learned from data and the equations generally differ from the epistatic weights given as in (4), but similar biological and geometric conclusions can be drawn. This idea could then be taken further by considering smoothing splines, instead of linear regression, and their relation to epistatic filtrations. From an application point of view, one would obtain an interesting new extension of the concept of epistasis because intermediate genotypes could be assessed, which would correspond to the case of genetically heterogeneous populations of organisms as occur in nature.

Other numerical results for the above regressions are summarized in Table S9. Over 0**** two coefficients are significantly non-zero (for *x*_1_ and *x*_4_), see top part of Table S9. Similarly, over 1**** four coefficients are significantly non-zero (*x*_1_, *x*_2_, *x*_3_, *x*_4_), see bottom part of Table S9. The fit of the linear regression models is confirmed by the relatively high values of the *F*-statistic. Over 0\*\*\**v* the *F*-statistics is 459.1 for a *p*-value near zero and 720 observations. Over 1**** the corresponding *F*-statistics (near zero) is 52.61.

### B.13 Comparison with other approaches

Currently the main lines of research to investigate higher-order epistasis in computational biology and related disciplines include the present methods, inspired from discrete polyhedral geometry (*3, 33, 38, 62*); linear regression approaches, c.f. (*21*); methods originating from harmonic analysis, c.f. (*40, 54, 63*); and using correlations between the effects of pairwise mutations, discussed in (*38*).

In a 2-locus, biallelic system, all these methods can easily be recovered from one another; some of them even agree. This is true also for some ecological approaches, including the generalized Lotka-Voleterra equations, which yield a mathematically equivalent form to epistasis for certain situations c.f. see equation 9 of (*4*). In higher dimensional systems, these methods remain conceptually closely related but they generally yield different insights about the problem, such as whether the interactions are significant, what their magnitude is, and what their sign is. Because these previous methods make specific, *a priori* assumptions about the forms of interactions, they are limited by these assumptions. Epistatic filtrations add a global perspective, determining the structure of interactions from the shape of the fitness landscape.

### B.14 Microbiome data sets

In this work, *Drosophila* microbiome fitness landscapes consist of experimental measurements on germ-free *Drosophila* flies inoculated with different bacterial species. The lifespan of approximately 100 individual flies were measured for each combination of bacterial species, giving roughly 3,200 individual fly lifespans for each of the two data sets presented. The experimental methods are described in (*3, 64*). The first data set is the exact data presented in (*3, 64*). The second data set is the second set of species with exactly the same methods used in (*3, 64*). The bacterial compositions considered consist of all possible combinations of five species. The species considered can all occur naturally in the gut of wild flies: *Lactobacillus plantarum* (LP), *Lactobacillus brevis* (LB), *Acetobacter pasteurianus* (APa), *Acetobacter tropicalis* (AT), *Acetobacter orientalis* (AO), *Acetobacter cerevisiae* (AC), *Acetobacter malorum* (AM). The 5-member communities both stably persist in the fly gut. For the purposes of this work, we define **stable** as maintaining colonization of the gut when ≤ 20 flies are co-housed in a standard fly vial and transferred daily to fresh food containing 10% glucose, 5% live yeast that has subsequently been autoclaved, 1.2% agar, and 0.42% propionic acid, with a pH of 4.5. The total number of species found stably associated with an individual fly is typically between 3 and 8. Consistently, *Lactobacillus plantarum* and *Lactobacillus brevis*, are found with two to three *Acetobacter* species. Less consistently, species of *Enterobacteria* and *Enterococci* occur, and these have been described as pathogens. While more strains may be present, for each of the two data sets in the present work, a set of five non pathogen species was chosen, including the two *Lactobacilli* and three *Acetobacter* species. The combinations of species are shown in Table S7. Different strains of the same species were used in the two data sets.

---

**Figure S1:**
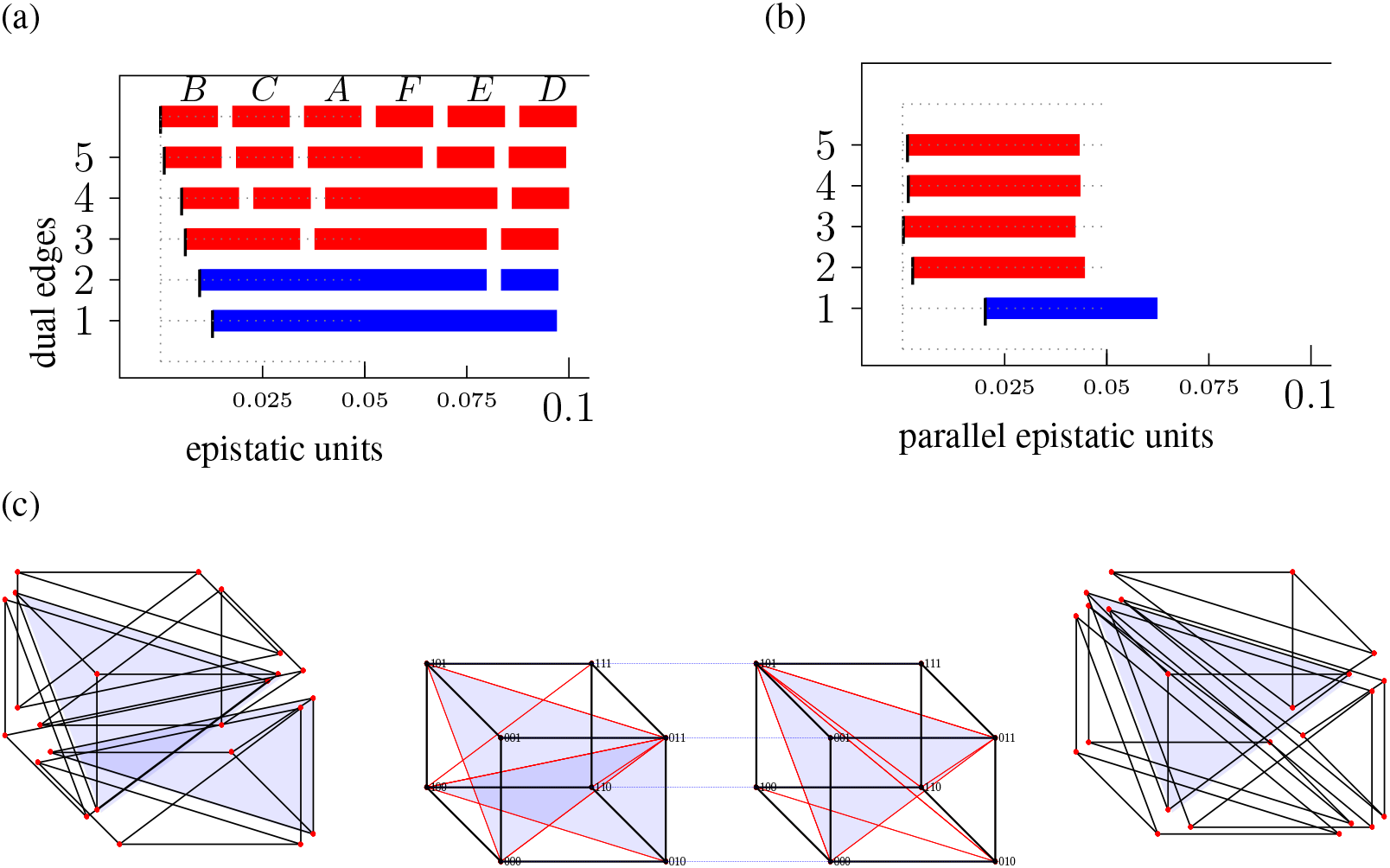
Parallel transport from 0**0* to 1**0* within the Khan dataset. (a) Filtration based on the triangulation of 0**0*. (b) Parallel epistatic weights computed from 1**0* for the triangulation based on 0**0*. (c) The two parallel triangulations (and exploded copies) are depicted. The partitions in the node set are transferred from the cube on the middle left to the cube on the middle right. Exploded versions of these same triangulation on the far left and far right demonstrate the geometry of the simplices generated by the triangulations.

**Figure S2:**
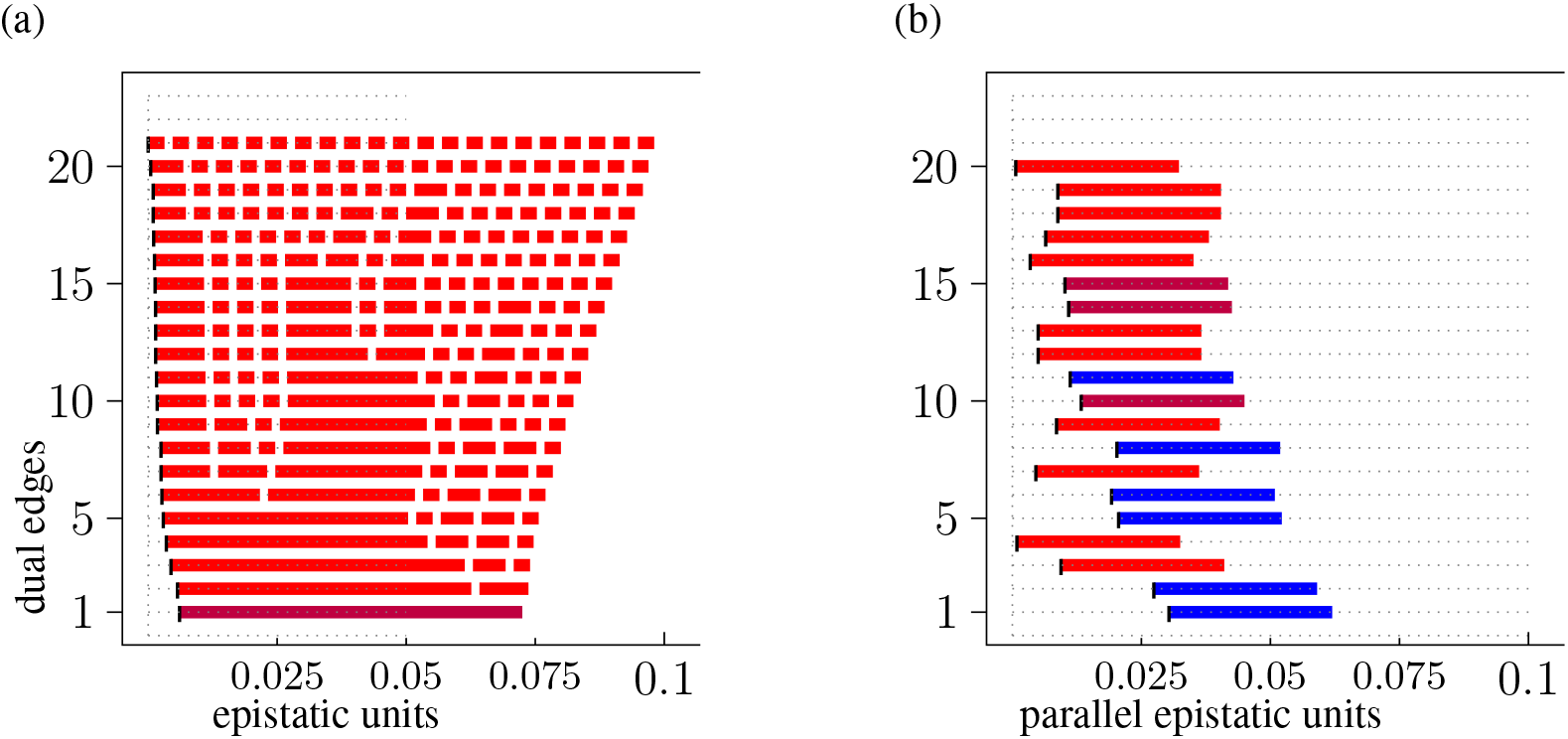
Epistatic filtration and parallel epistatic units for transport from ****0 to ****1 within the Khan data.

**Figure S3:**
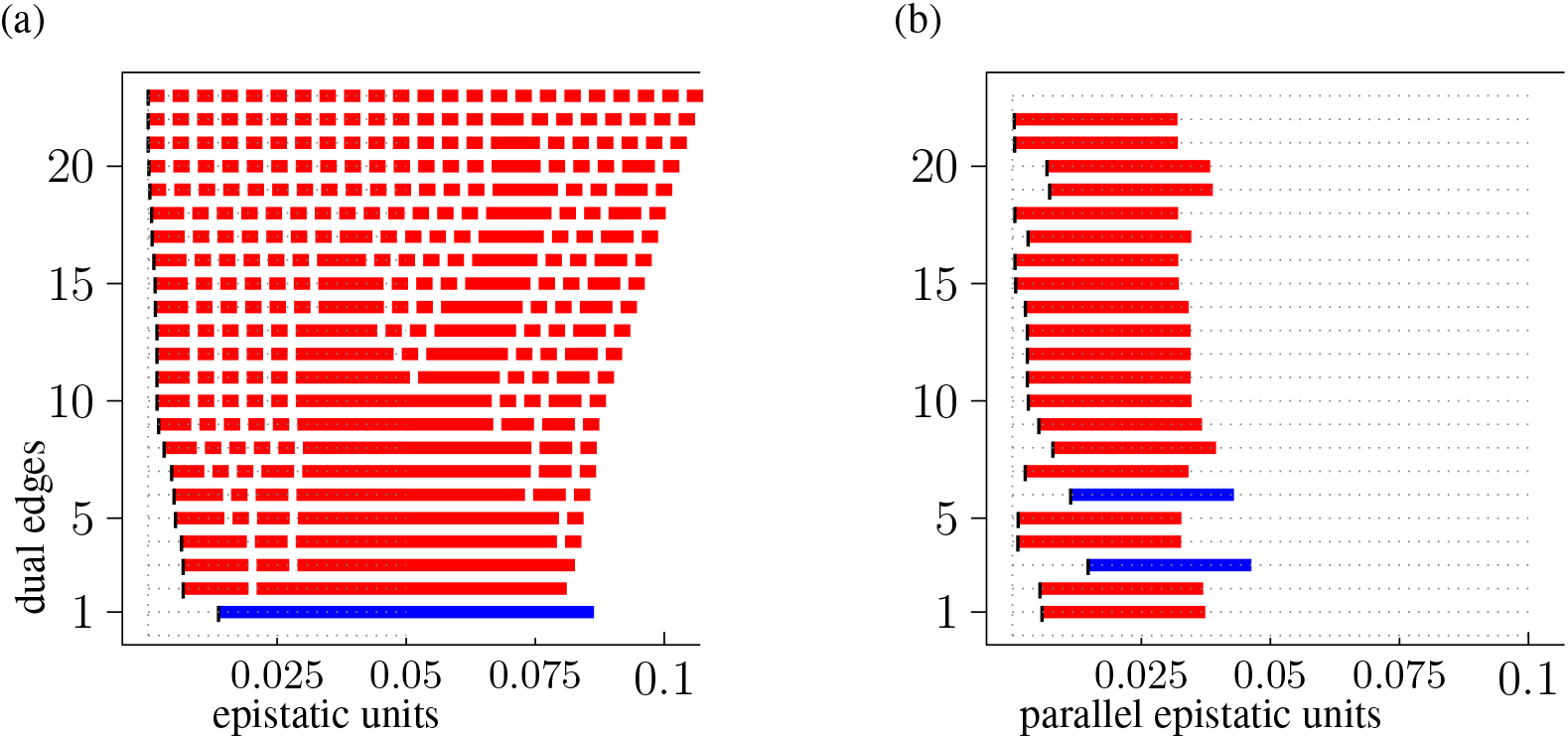
Epistatic filtration and parallel epistatic units for transport from ****1 to ****0 within the Khan data.

**Figure S4:**
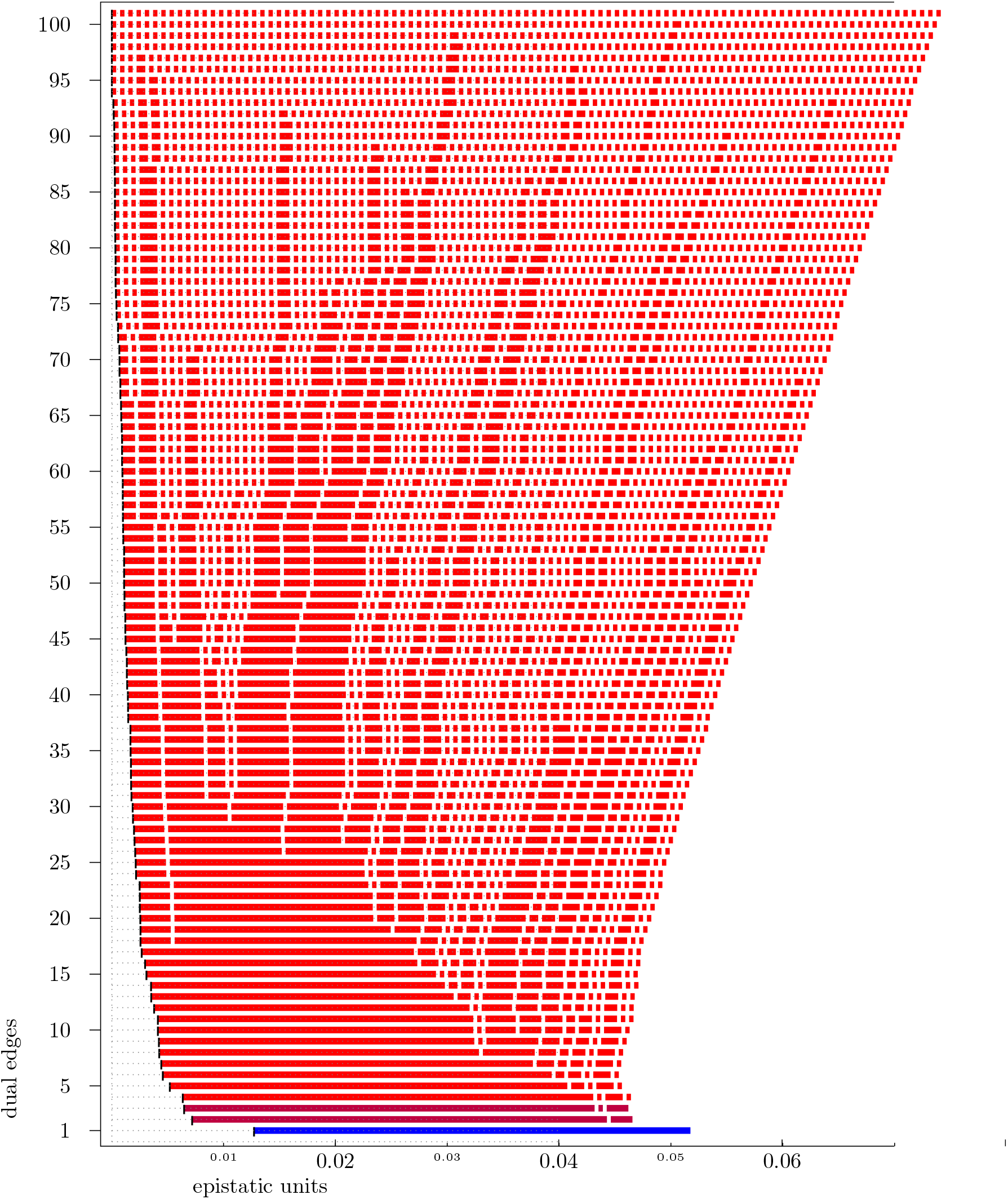
Complete filtration of the Khan data over the whole 5-cube.

**Figure S5:**
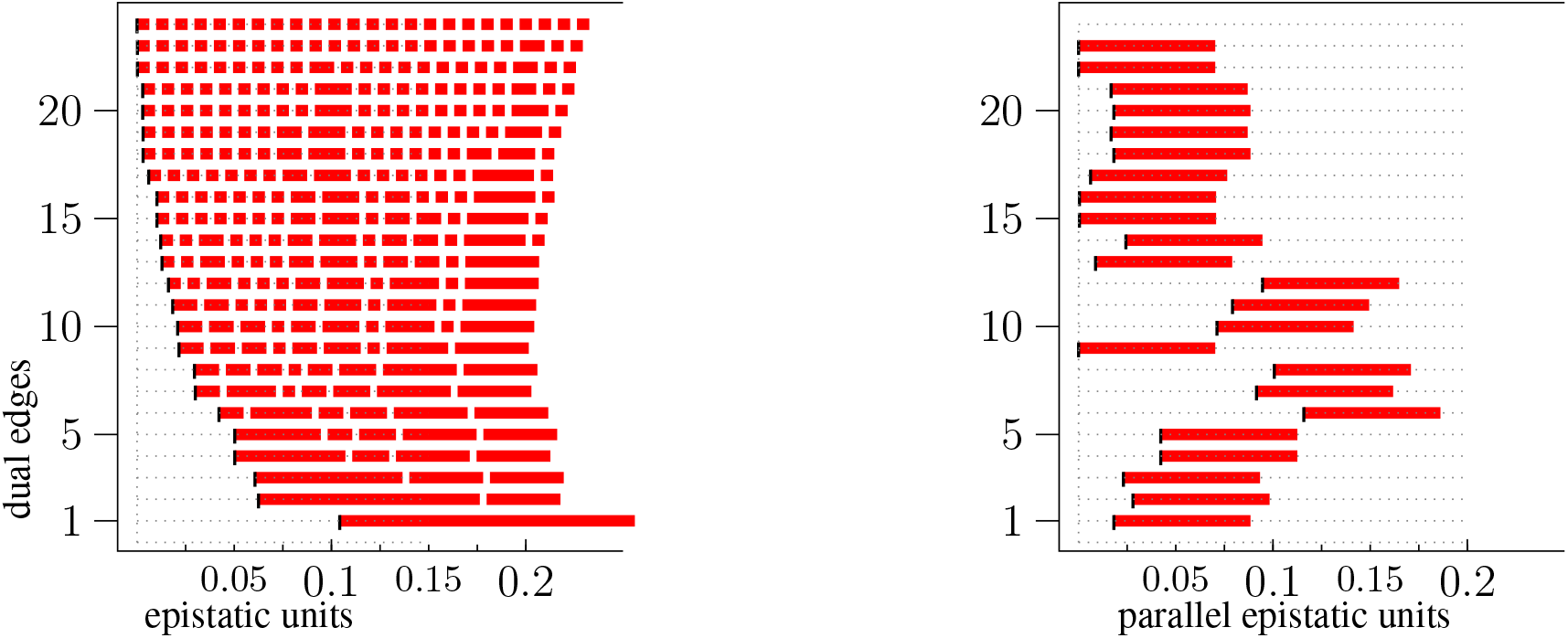
Parallel transport from 0**** to 1**** within the Tan data. Analysis based on mean values only; hence there is no color coding for the significance.

**Figure S6:**
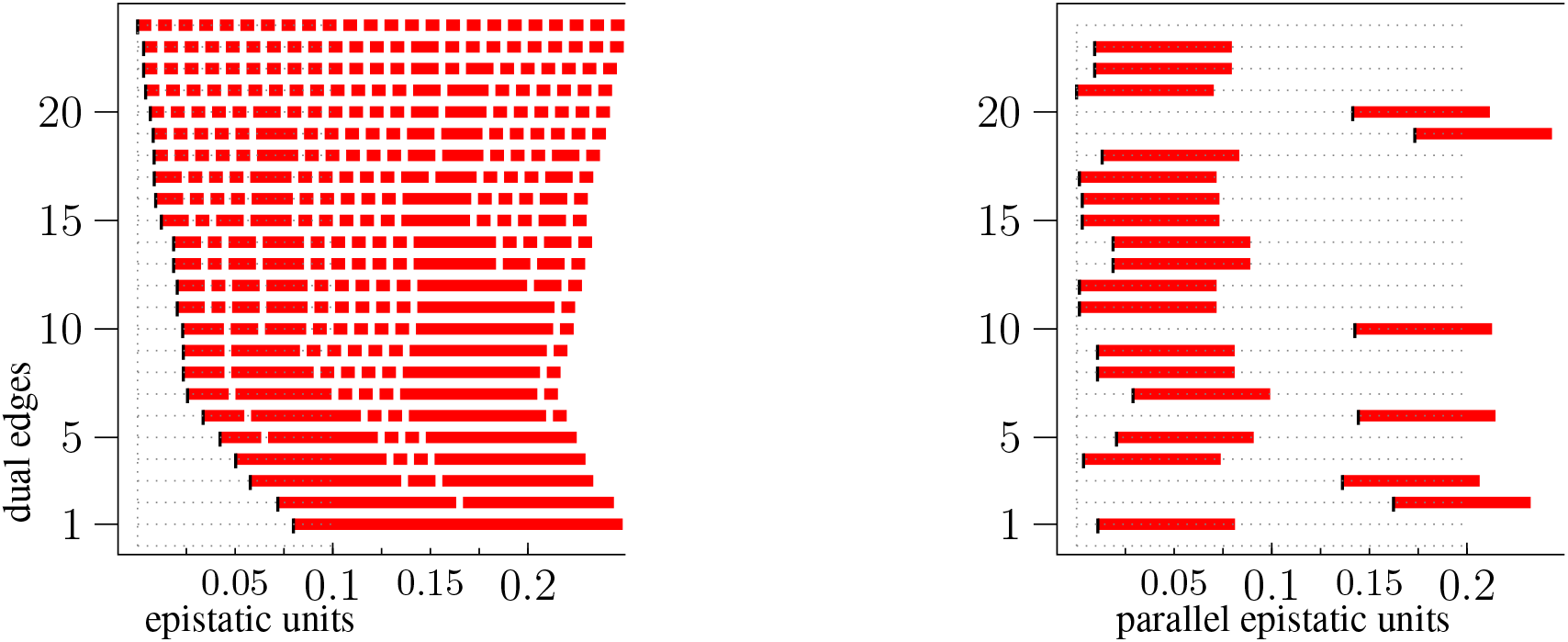
Parallel transport from the face **0* to the face **1** within the Tan data. Analysis based on mean values only; hence there is no color coding for the significance.

**Figure S7:**
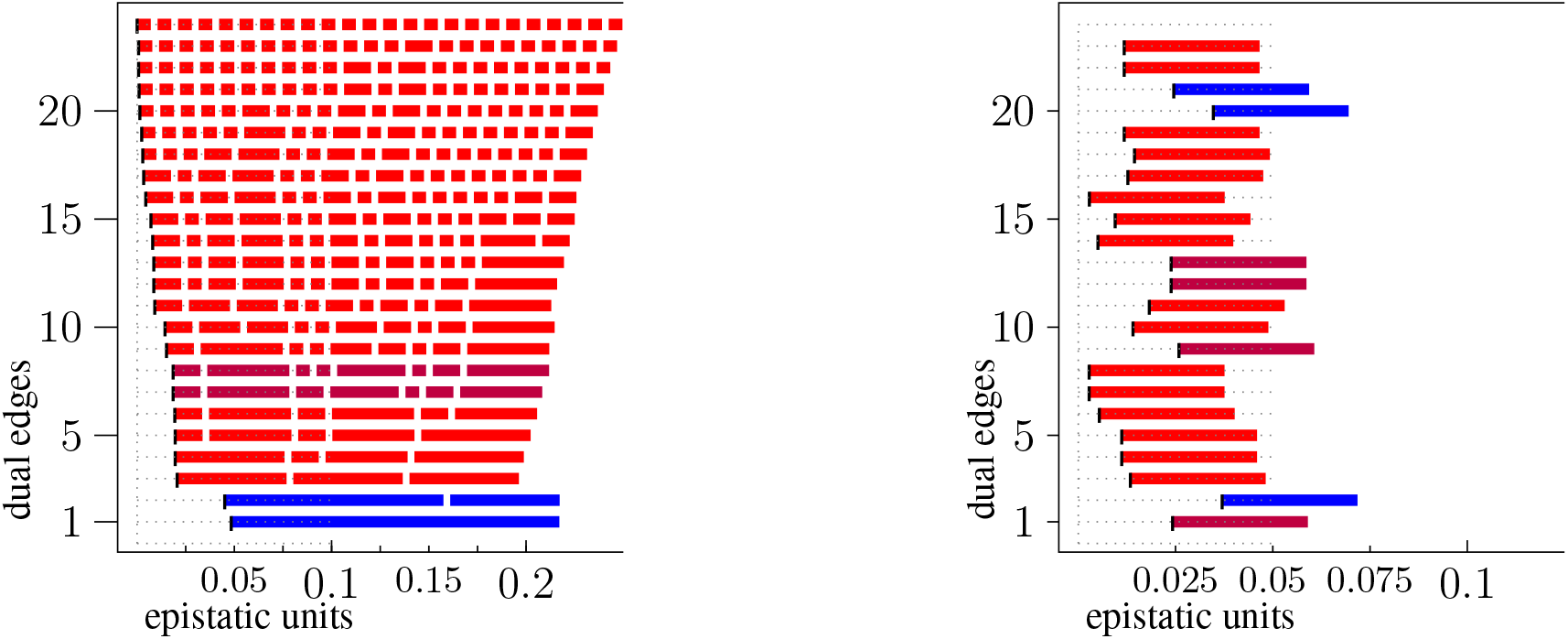
0****(Eble) to 1****(Eble).

**Figure S8:**
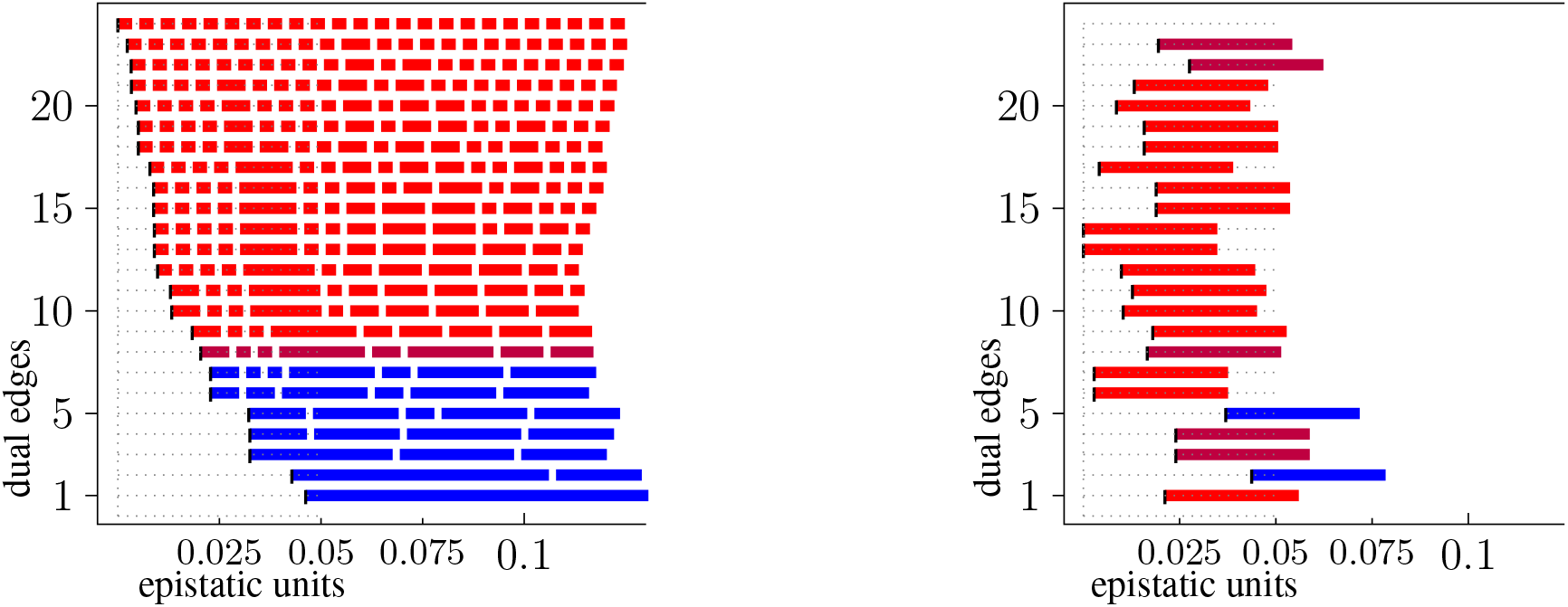
*0***(Eble) to *1***(Eble).

**Figure S9:**
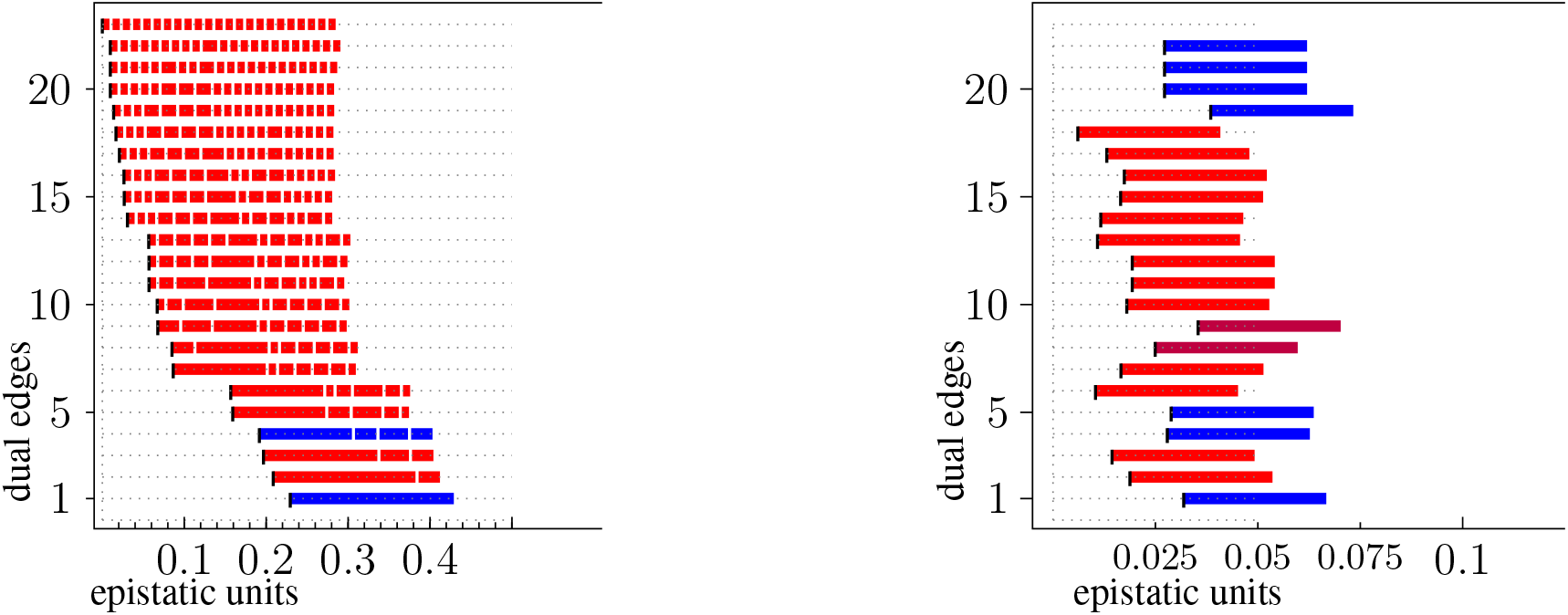
0****(GouldCFU) to 0****(GouldTTD).

**Figure S10:**
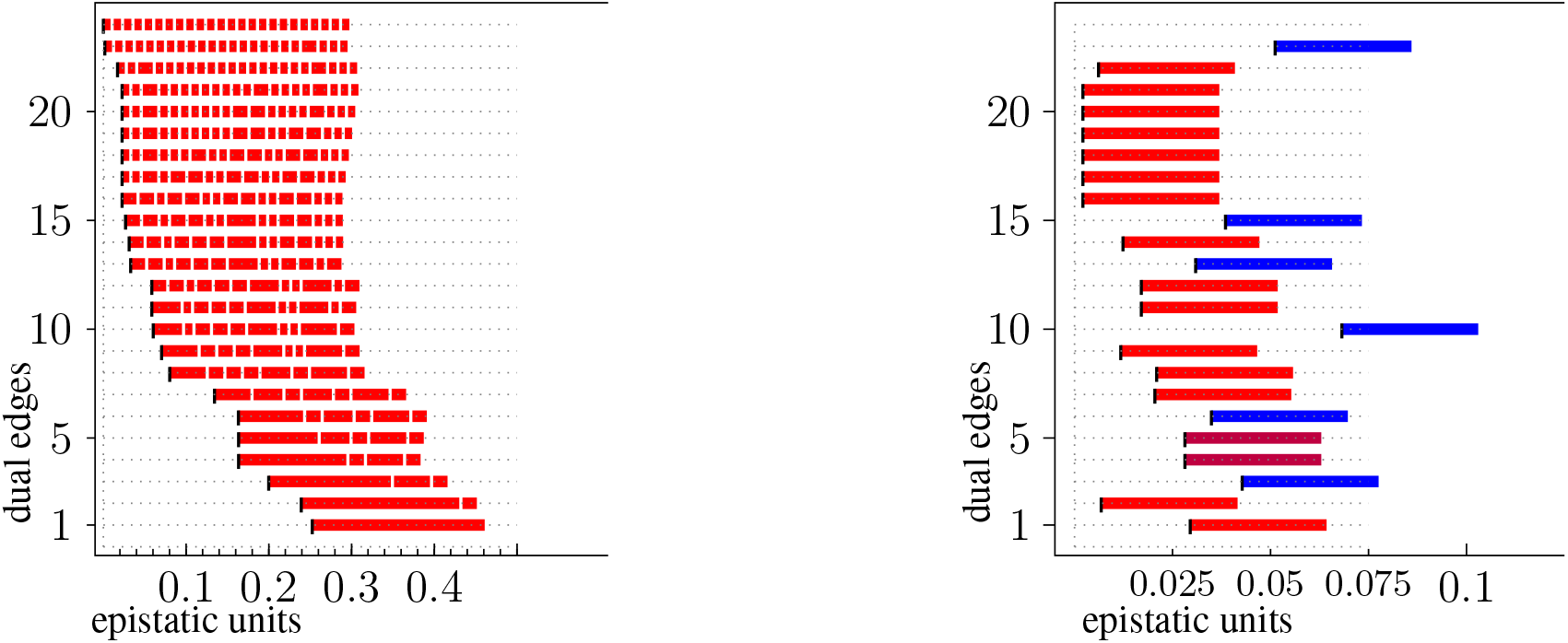
1****(GouldCFU) to 1****(GouldTTD).

**Figure S11:**
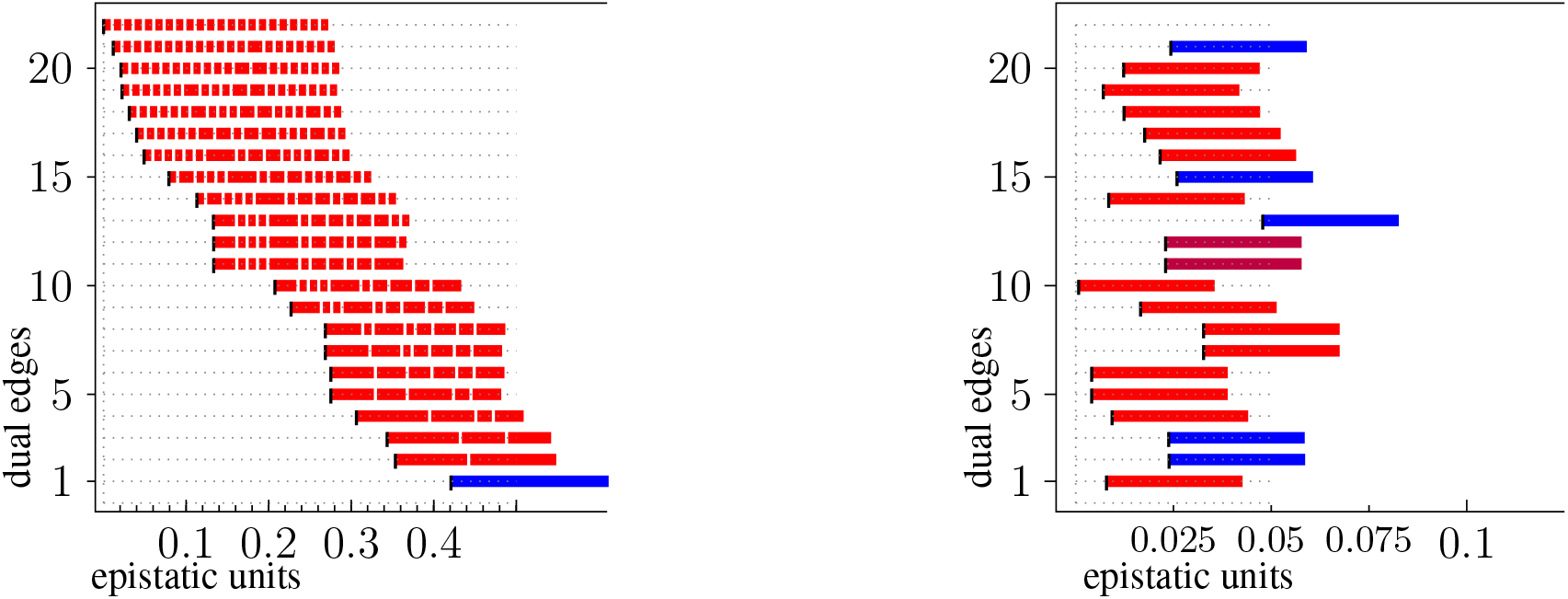
*0***(GouldCFU) to *0***(GouldTTD).

**Figure S12:**
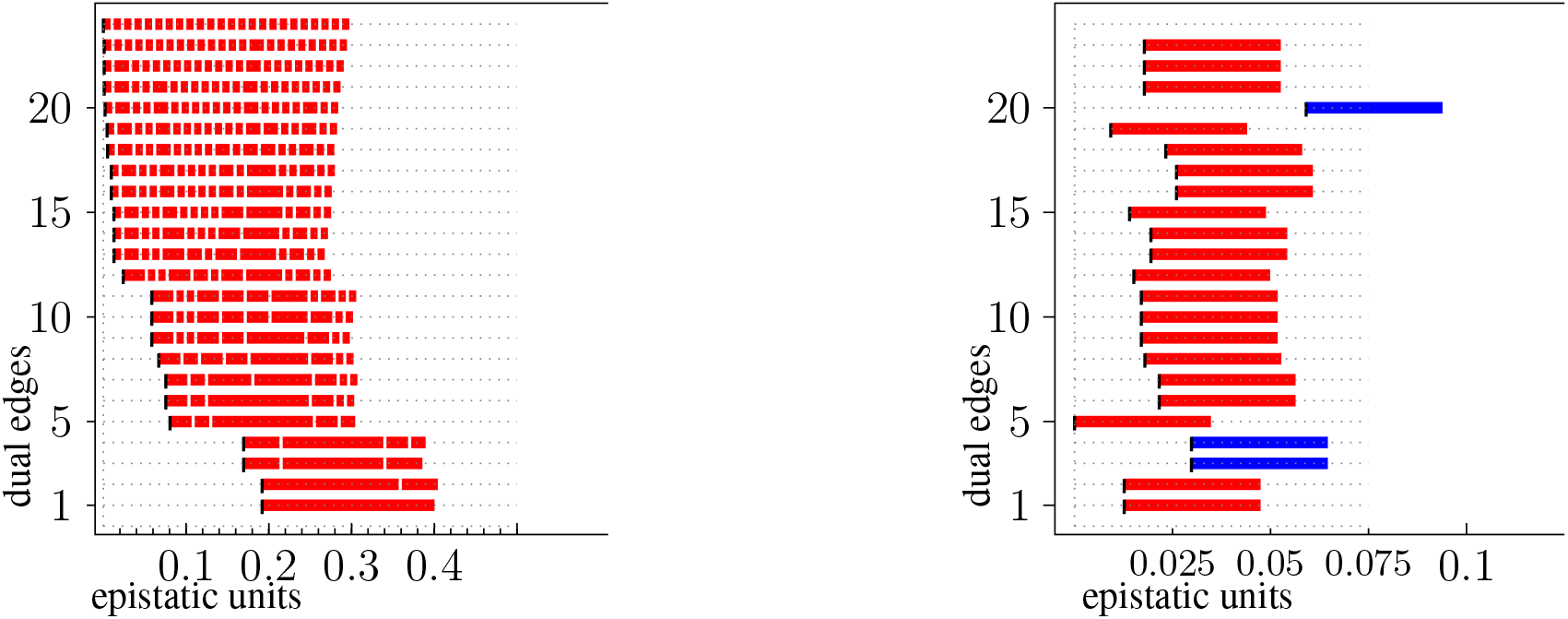
*1***(GouldCFU) to *1***(GouldTTD).

**Figure S13:**
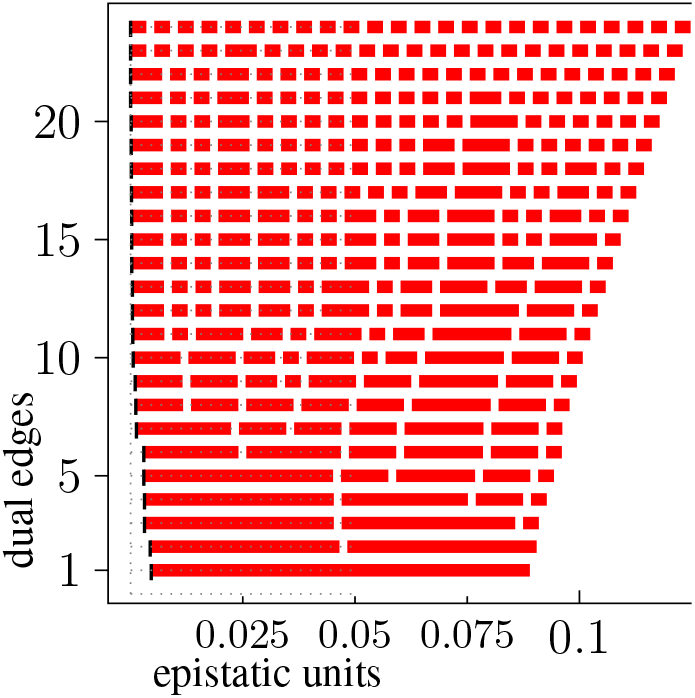
Product model associated to the parallel transport 0**0* → 1**0* within the Khan evolution data, cf. (Fig. S1.)

**Figure S14:**
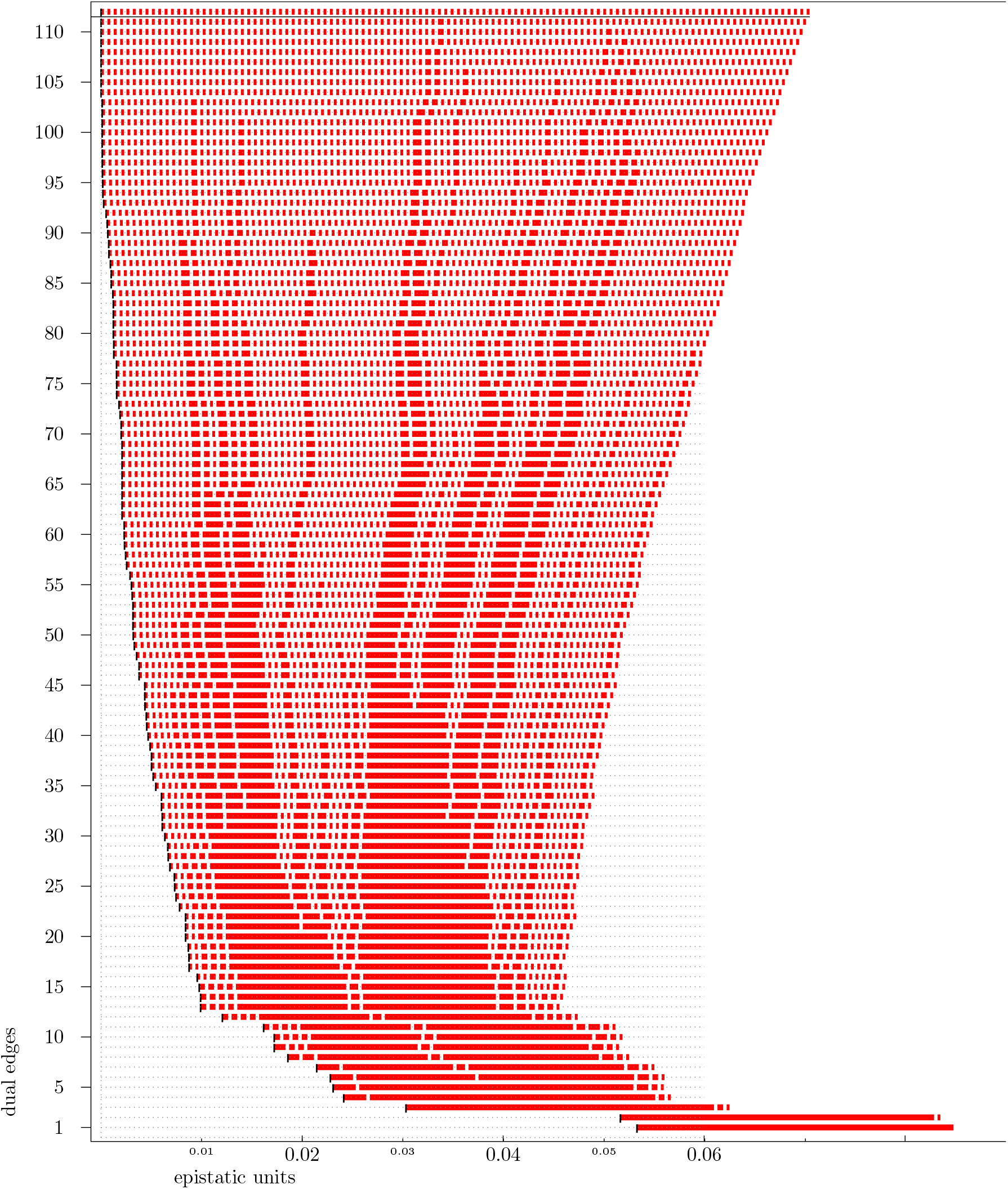
Non-generic product model associated to the parallel transport **0** → **1** within the Tan data. Its unique non-simplicial maximal cell has 7 vertices and is split into a bipyramid by a slight perturbation of its height values, cf. Theorem 8 of (*38*). The corresponding artificial dual edge has edge label 111 and is indicated by a horizontal line.

**Figure S15:**
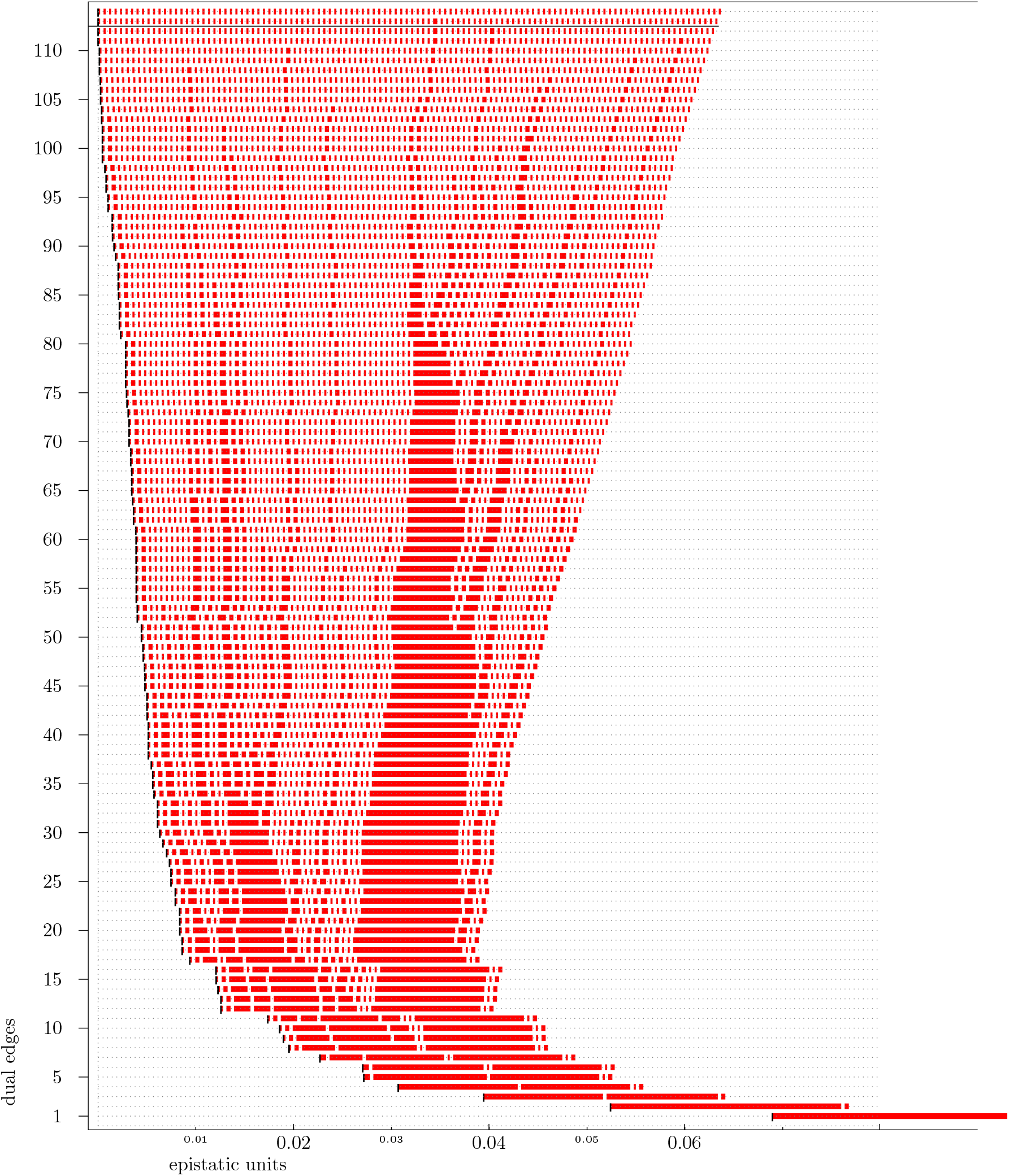
Non-generic product model associated to the parallel transport 0**** → 1**** within the Tan data. There are two non-simplicial maximal cells, both of cardinality 7. As in (Fig. S14) they are split into a bipyramid each at the beginning of the filtration process.

**Figure S16:**
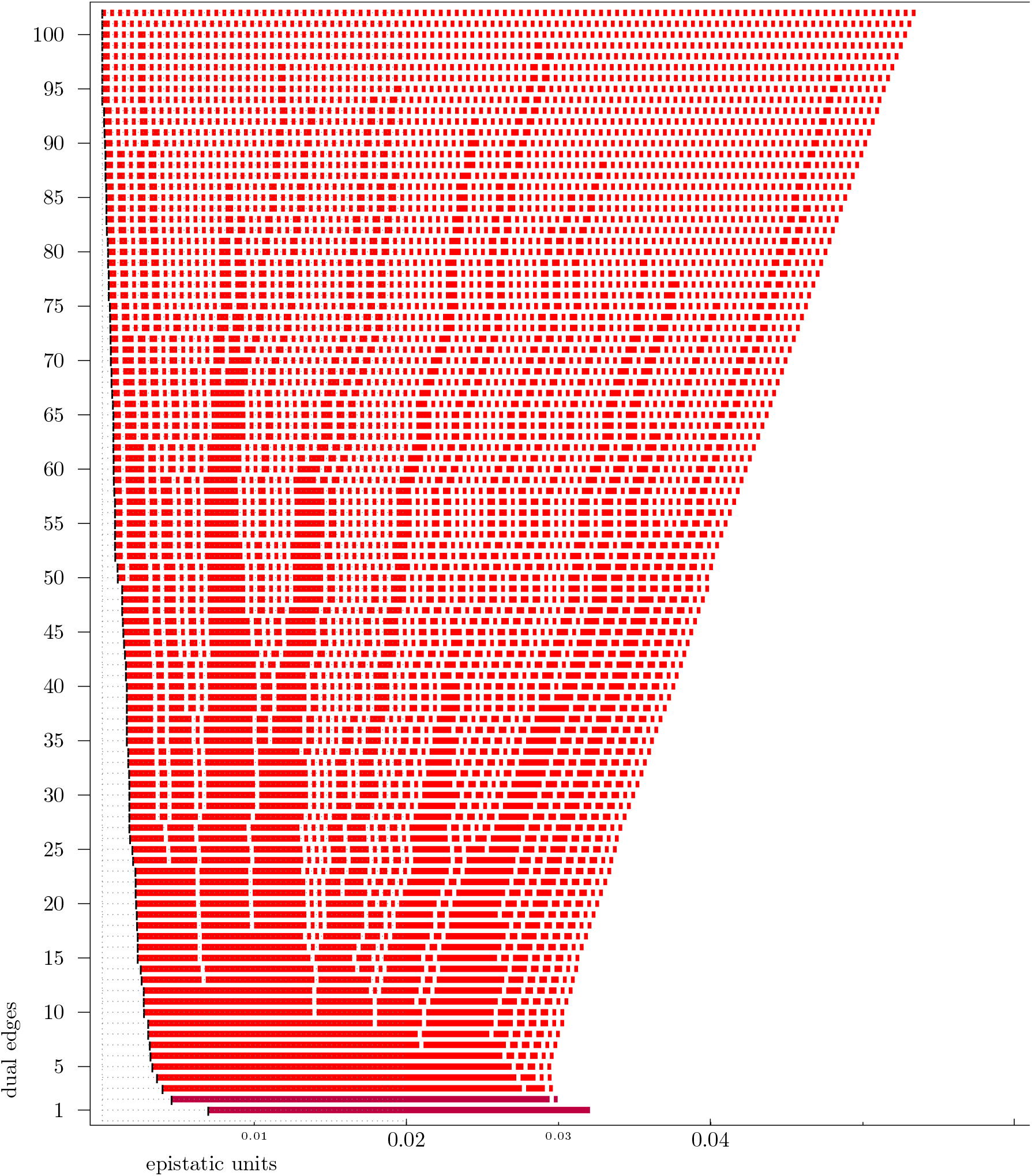
Product model for the parallel transport Khan ***0* → ***1*. The semisignificant bipyramid labeled 2 reads {(1000)_*o*_} + {(0000)_*o*_, (1010)_0_, (0110)_*o*_, (1010)_*p*_, (0011)_*p*_} + {(0010)_*o*_} and the semisignificant bipyramid labeled 1 reads {(1100)_*o*_} + {(1011)_*o*_, (1010)_*p*_, (1001)_*p*_, (0011)_*p*_, (1111)_*p*_} + {(1011)_*p*_}.

**Figure S17:**
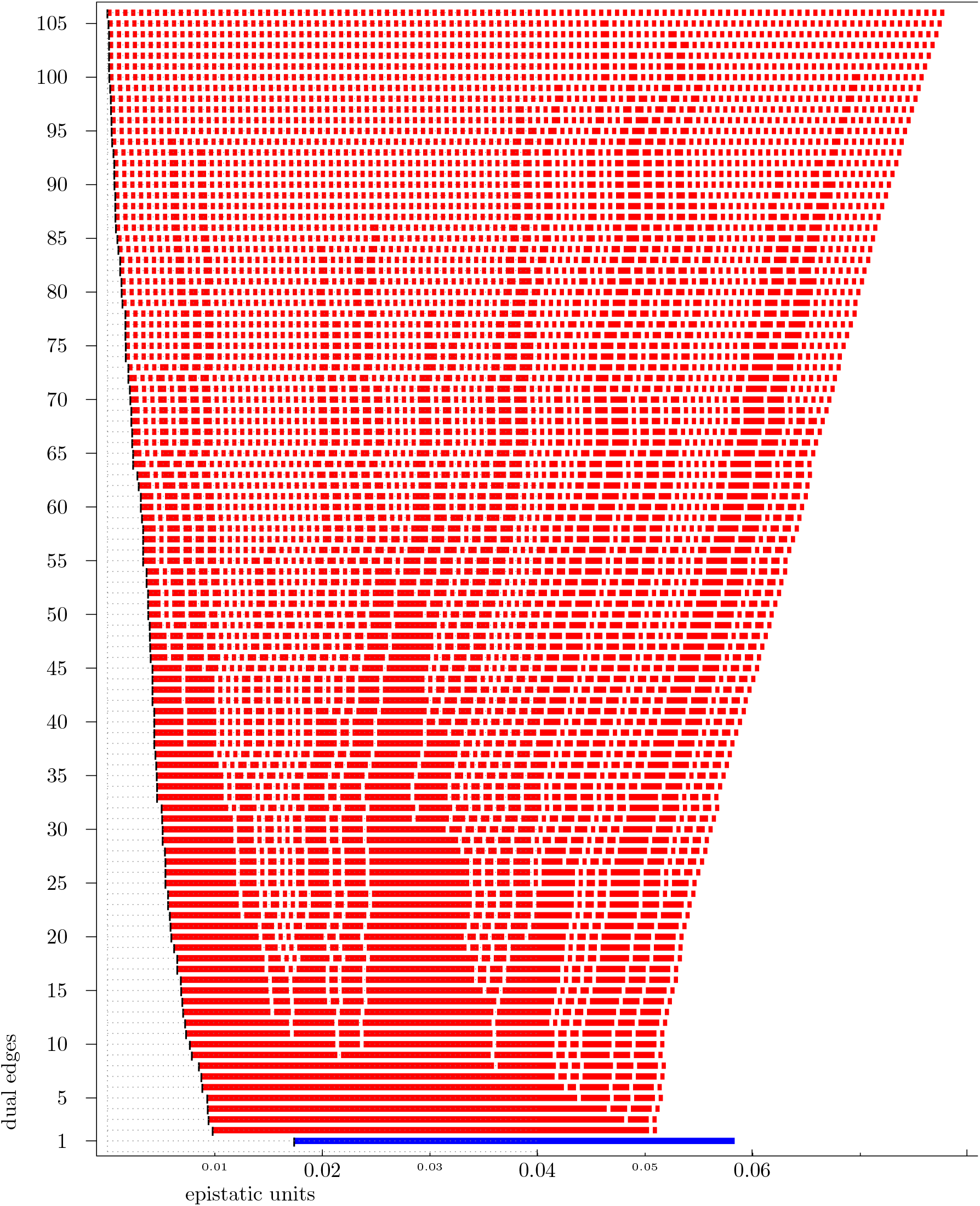
Product model for the parallel transport Eble 0**** → 1****. The unique significant bipyramid reads {(0001)_*o*_} + {(0000)_*o*_, (1001)_*o*_, (0101)_*o*_, (0011)_*o*_, (0001)_*p*_} + {(0101)_*p*_}.

**Figure S18:**
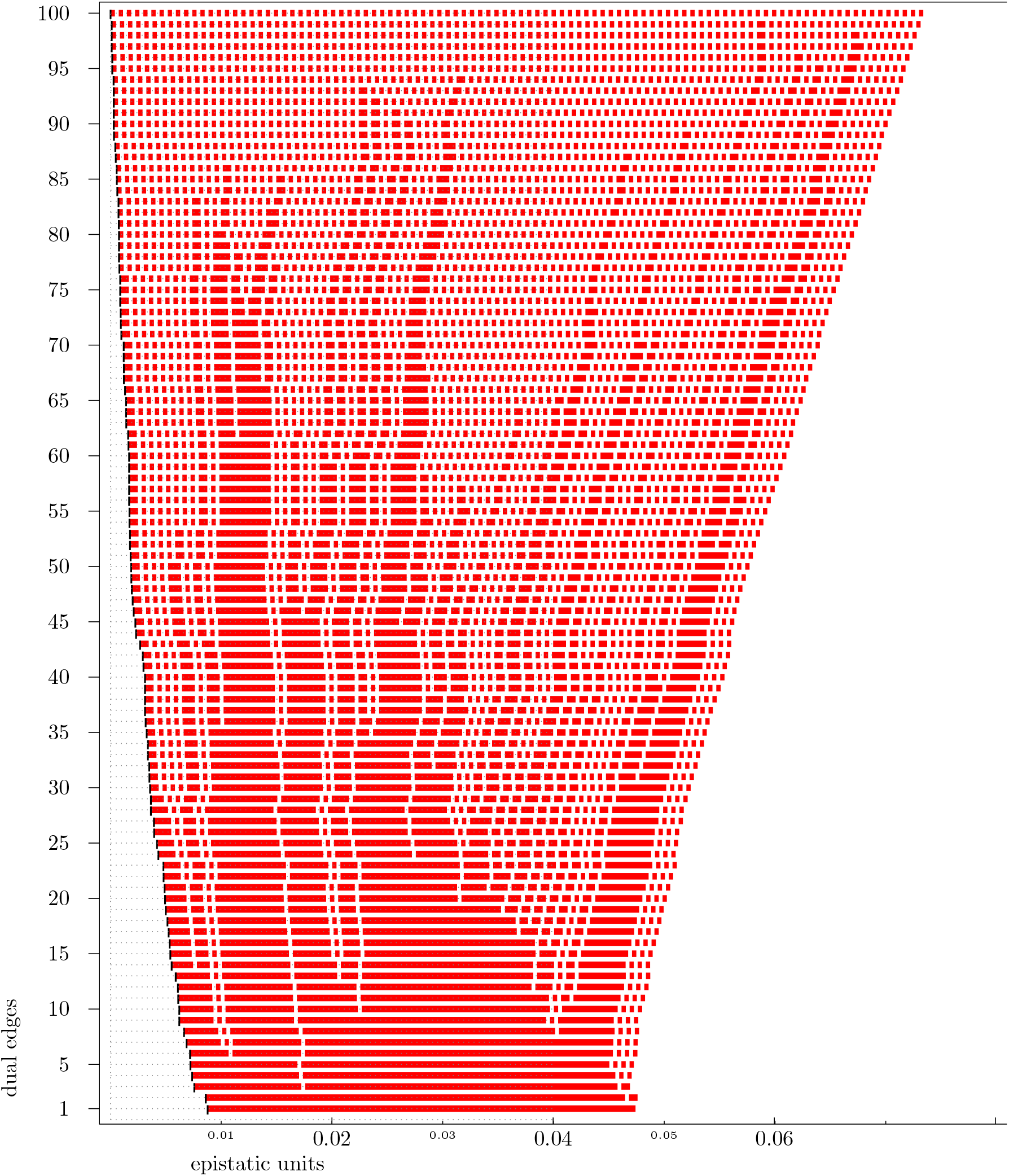
Product model for the parallel transport Eble *0*** → *1***.

**Figure S19:**
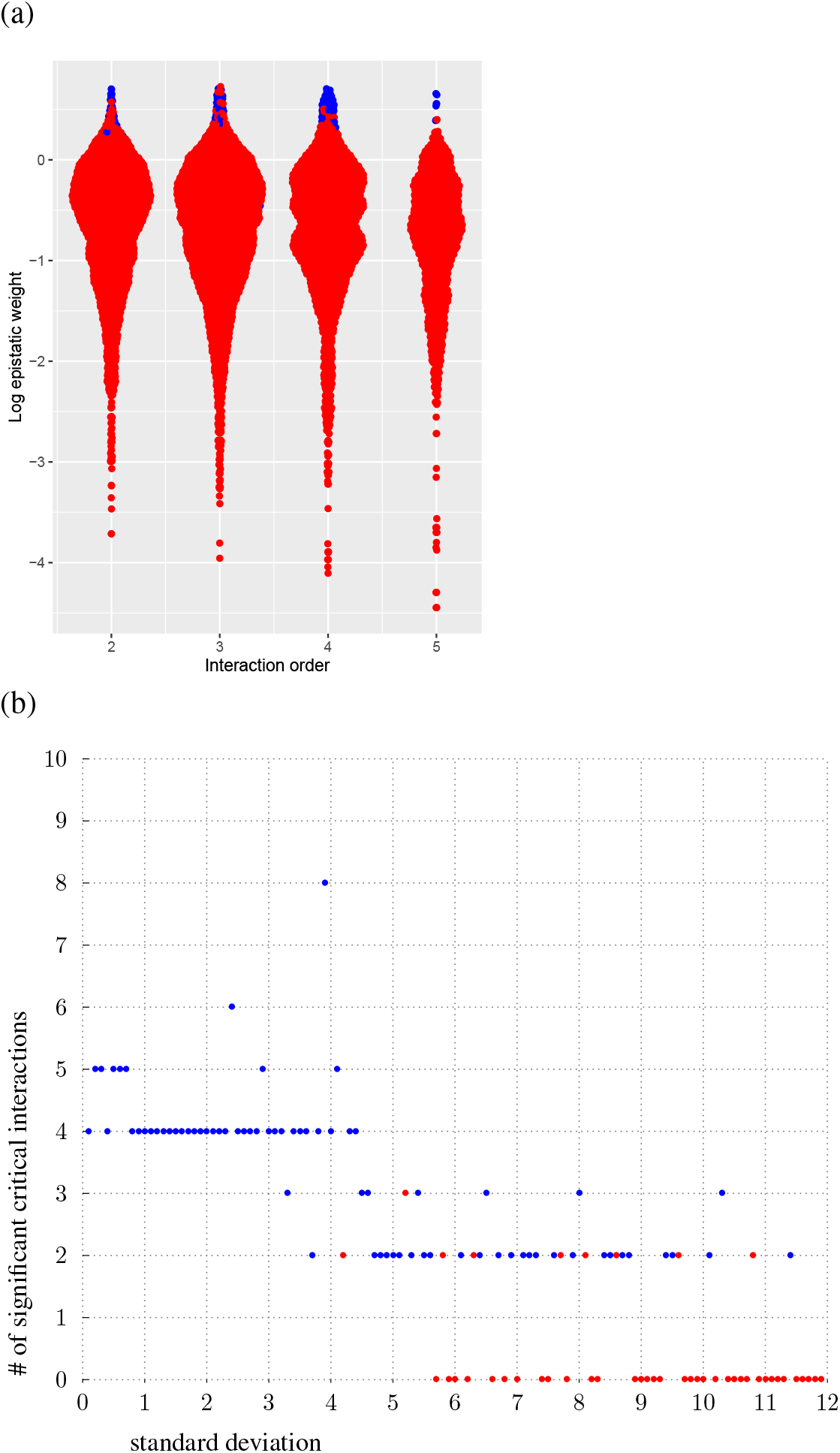
Synthetic data demonstrate method performance. Synthetic height functions over the 4-dimensional cube are generated with 100 replicates each and standard deviation as indicated. The heights of the wild type 0000 and 0001 are sampled with mean 53, all the other vertices with mean 50. (a) The distribution of log_10_-transformed epistatic weights is roughly constant as a function of interaction order, indicating the dimensional normalization is effective. (b) The number of significant interactions decreases as the standard deviation of the input data for each genotype increases. A blue dot is drawn if the interaction is significant and a red dot is drawn otherwise.

**Figure S20:**
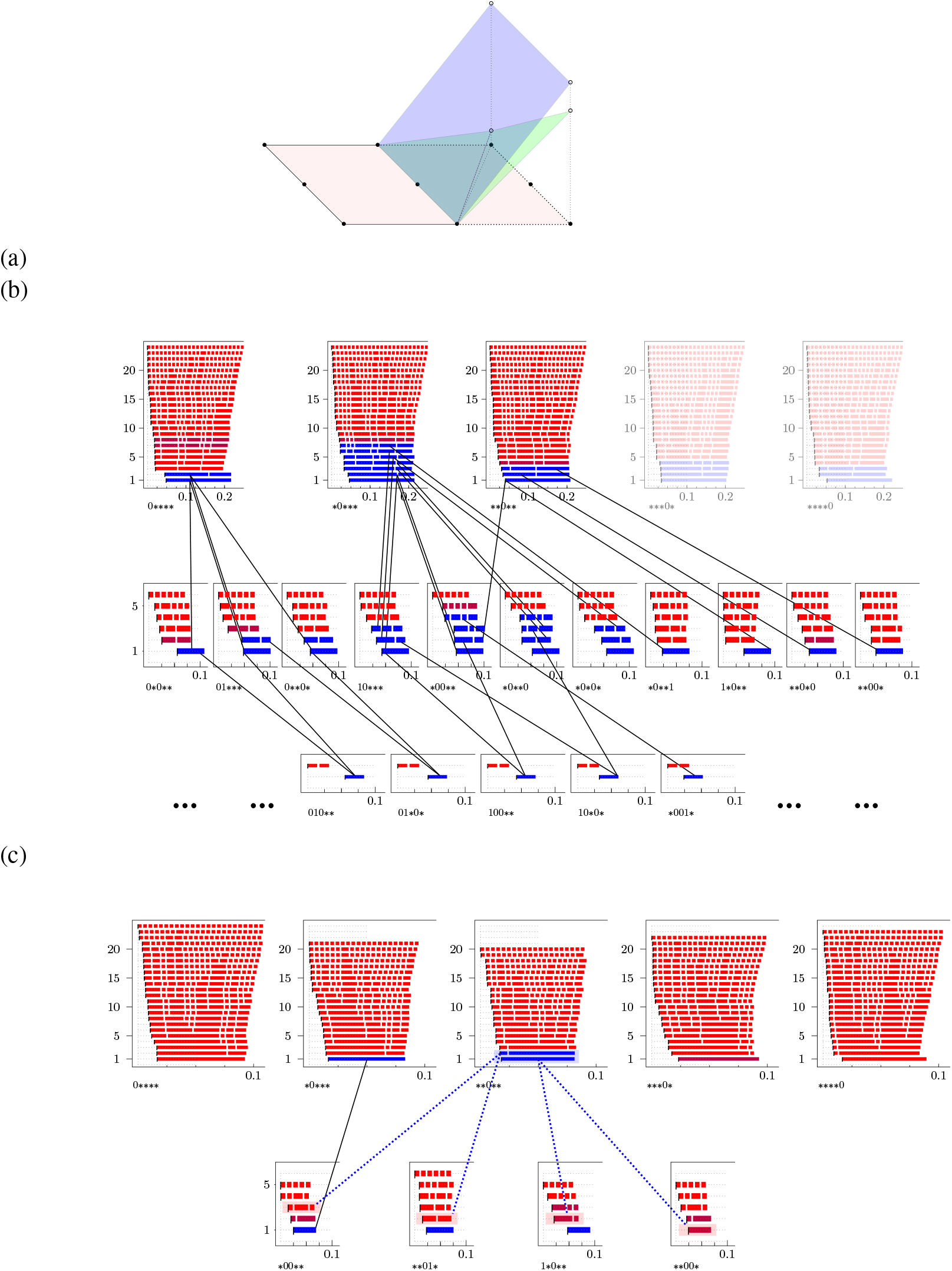
Meta-epistatic charts illustrate whether or not higher-order interactions arise from lower-order interactions. (a) Cartoon of the principle underlying meta-epistatic charts. The important loci in the interaction are depicted as black dots in a hyperplane through the genotypes, where the true dimensions of the genotypes are flattened onto the cartoon plane (pink). Higher-order interactions that derive from lower-order interactions occur in a new hyperplane (blue), which magnifies the weights of a subset of the landscape. In contrast, novel higher-order interactions that only arise in higher dimensions do not lie in a single additional hyperplane but instead require at least two additional hyperplanes (green). In (b) and (c) two meta-epistatic charts are represented. In each chart we identify the source of a higher-order interaction for the Eble and Gould data respectively. The results are compiled in Table S6.

**Figure S21:**
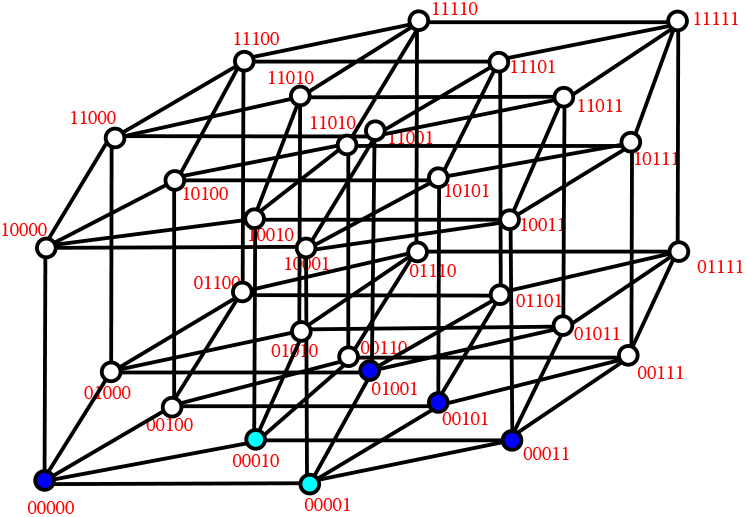
Vertices of the bipyramid {00001} + {00000, 01001, 00101, 00011} + {00010} arising for the Khan data set (*37*) restricted to *n* = 4 loci. Dark blue dots correspond to common face *s ∩ t* of the bipyramid and light blue dots correspond to the satellite vertices of *s* and *t*.

**Figure S22:**
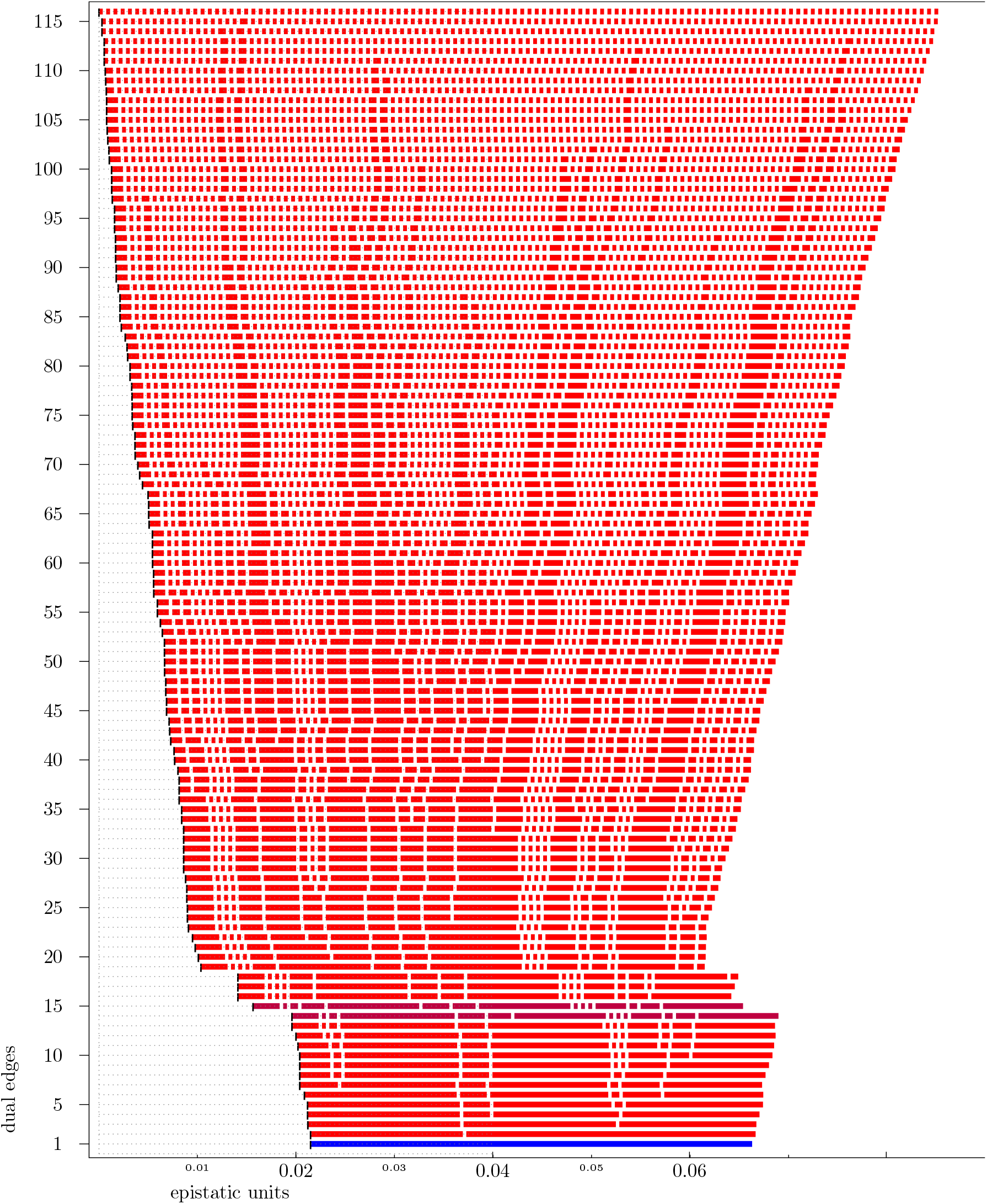
Complete filtration of the Eble fitness landscape over the whole 5-cube.

**Table S1:** Number of circuits of [0, 1]^*n*^ and bipyramids among these.

| dimensions | circuits | bipyramids | percentage |
| --- | --- | --- | --- |
| 2 | 1 | 1 | 100.00% |
| 3 | 20 | 8 | 40.00% |
| 4 | 1348 | 1088 | 80.71% |
| 5 | 353616 | 309056 | 87.40% |

**Table S2:**
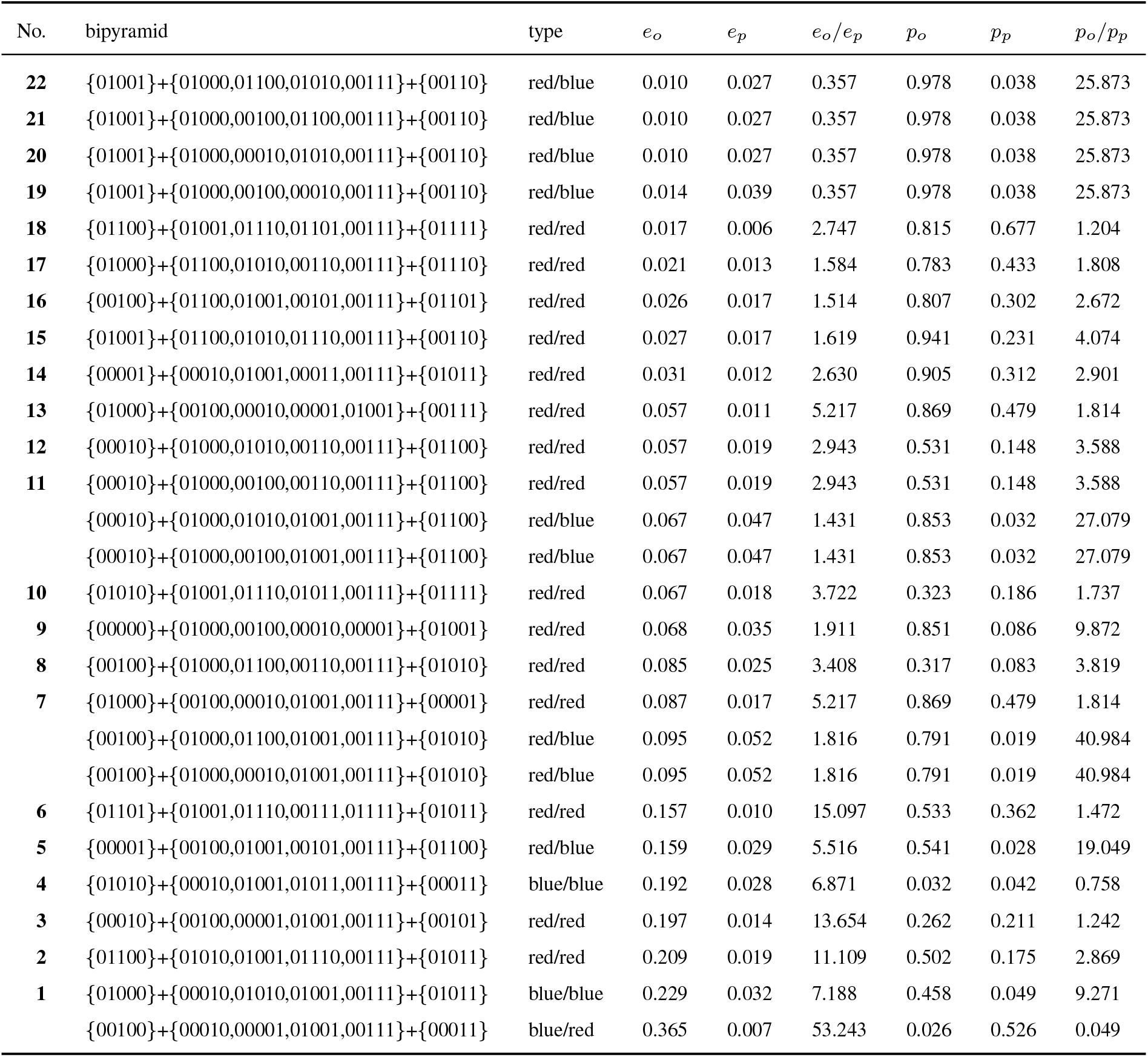
Parallel analysis GouldCFU 0**** → Gould 0****, non-critical red/red-case omitted.

**Table S3:**
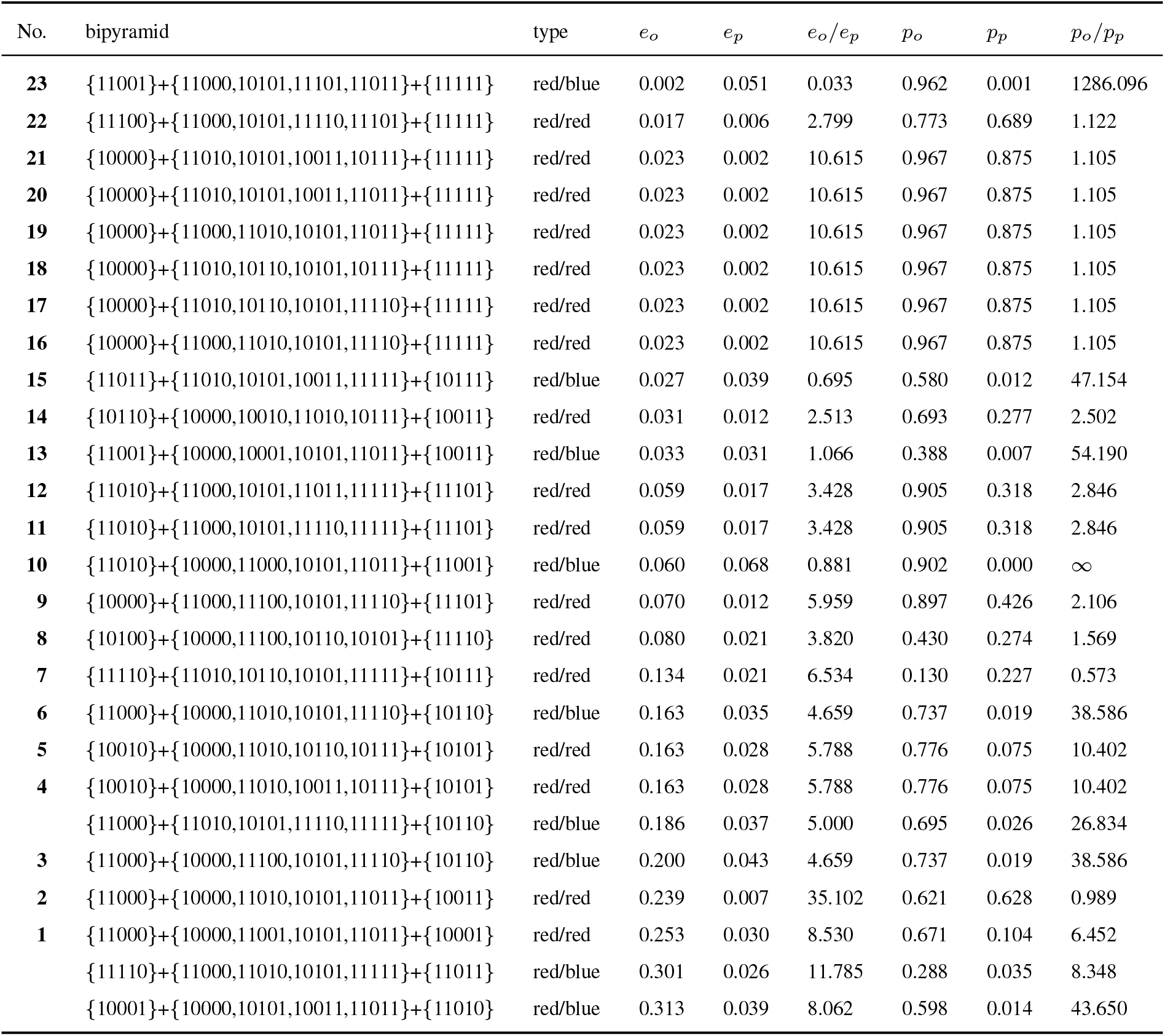
Parallel analysis GouldCFU 1**** → Gould 1****, non-critical red/red-case omitted.

**Table S4:** Parallel analysis GouldCFU *0*** → Gould *0***, non-critical red/red-case omitted.

| No. | bipyramid | type | $e_o$ | $e_p$ | $e_o/e_p$ | $p_o$ | $p_p$ | $p_o/p_p$ |
| --- | --- | --- | --- | --- | --- | --- | --- | --- |
| 21 | {10001}+{10000,00001,10101,10011}+{00111} | red/blue | 0.012 | 0.024 | 0.481 | 0.963 | 0.026 | 36.756 |
| 20 | {00010}+{10000,10010,00011,00111}+{10011} | red/red | 0.021 | 0.012 | 1.714 | 0.797 | 0.270 | 2.952 |
| 19 | {10100}+{10000,00100,10110,10101}+{00110} | red/red | 0.022 | 0.007 | 3.155 | 0.869 | 0.717 | 1.212 |
| 18 | {10110}+{10000,10010,00111,10111}+{10011} | red/red | 0.031 | 0.012 | 2.513 | 0.693 | 0.277 | 2.502 |
| 17 | {00100}+{10000,00110,10101,00111}+{10110} | red/red | 0.040 | 0.018 | 2.266 | 0.915 | 0.290 | 3.155 |
| 16 | {00100}+{10000,00110,10110,10101}+{00111} | red/red | 0.049 | 0.022 | 2.266 | 0.915 | 0.290 | 3.155 |
| 15 | {00001}+{10000,00100,00010,00111}+{00110} | red/blue | 0.079 | 0.026 | 3.047 | 0.698 | 0.023 | 30.749 |
| 14 | {00010}+{10000,10010,00110,00111}+{10110} | red/red | 0.113 | 0.008 | 13.352 | 0.295 | 0.461 | 0.640 |
| 13 | {00000}+{10000,00100,00010,00001}+{00111} | red/blue | 0.133 | 0.048 | 2.775 | 0.476 | 0.001 | 707.281 |
| 12 | {10010}+{10000,10011,00111,10111}+{10101} | red/red | 0.133 | 0.023 | 5.788 | 0.776 | 0.075 | 10.402 |
| 11 | {10010}+{10000,10110,00111,10111}+{10101} | red/red | 0.133 | 0.023 | 5.788 | 0.776 | 0.075 | 10.402 |
| 10 | {00011}+{10000,00001,10011,00111}+{10101} | red/red | 0.208 | 0.001 | 275.689 | 0.413 | 0.949 | 0.435 |
| 9 | {00010}+{10000,00100,00001,00111}+{00101} | red/red | 0.227 | 0.017 | 13.654 | 0.262 | 0.211 | 1.242 |
| 8 | {00100}+{10000,00001,00101,00111}+{10101} | red/red | 0.269 | 0.033 | 8.193 | 0.579 | 0.101 | 5.733 |
| 7 | {00001}+{10000,00100,00101,00111}+{10101} | red/red | 0.269 | 0.033 | 8.193 | 0.579 | 0.101 | 5.733 |
| 6 | {00001}+{10000,00010,00011,00111}+{10010} | red/red | 0.275 | 0.004 | 67.167 | 0.493 | 0.755 | 0.653 |
| 5 | {00001}+{10000,00011,10011,00111}+{10010} | red/red | 0.275 | 0.004 | 67.167 | 0.493 | 0.755 | 0.653 |
| 4 | {00110}+{10000,00100,10101,00111}+{00101} | red/red | 0.306 | 0.009 | 32.813 | 0.413 | 0.610 | 0.677 |
| 3 | {00110}+{10000,10010,10110,00111}+{10111} | red/blue | 0.344 | 0.024 | 14.403 | 0.186 | 0.035 | 5.345 |
| 2 | {00110}+{10000,00010,10010,00111}+{00011} | red/blue | 0.354 | 0.024 | 14.760 | 0.175 | 0.030 | 5.853 |
|  | {00001}+{10000,10101,10011,00111}+{10111} | red/blue | 0.408 | 0.028 | 14.815 | 0.108 | 0.013 | 8.308 |
| 1 | {00100}+{10000,00010,00001,00111}+{00011} | blue/red | 0.421 | 0.008 | 53.243 | 0.026 | 0.526 | 0.049 |
|  | {00110}+{10000,10110,10101,00111}+{10111} | red/blue | 0.486 | 0.034 | 14.403 | 0.186 | 0.035 | 5.345 |

**Table S5:** Parallel analysis GouldCFU *1*** → Gould *1***, non-critical red/red-case omitted.

| No. | bipyramid | type | $e_o$ | $e_p$ | $e_o/e_p$ | $p_o$ | $p_p$ | $p_o/p_p$ |
| --- | --- | --- | --- | --- | --- | --- | --- | --- |
| 23 | {01100}+{11000,01110,01101,11110}+{11010} | red/red | 0.001 | 0.018 | 0.054 | 0.998 | 0.292 | 3.418 |
| 22 | {01100}+{11000,01010,01001,01110}+{11010} | red/red | 0.001 | 0.018 | 0.054 | 0.998 | 0.292 | 3.418 |
| 21 | {01100}+{11000,01001,01110,01101}+{11010} | red/red | 0.001 | 0.018 | 0.054 | 0.998 | 0.292 | 3.418 |
| 20 | {11001}+{11000,01001,11101,11011}+{11111} | red/blue | 0.002 | 0.059 | 0.033 | 0.962 | 0.001 | 1286.096 |
| 19 | {11110}+{11010,01110,01101,11111}+{01111} | red/red | 0.005 | 0.009 | 0.488 | 0.945 | 0.576 | 1.641 |
| 18 | {11100}+{11000,01100,11110,11101}+{01101} | red/red | 0.005 | 0.023 | 0.218 | 0.952 | 0.193 | 4.933 |
| 17 | {11000}+{11010,01110,01101,11110}+{11111} | red/red | 0.010 | 0.026 | 0.369 | 0.981 | 0.106 | 9.255 |
| 16 | {01110}+{11000,11010,01101,11110}+{11111} | red/red | 0.010 | 0.026 | 0.369 | 0.981 | 0.106 | 9.255 |
| 15 | {01100}+{11000,01101,11110,11101}+{11111} | red/red | 0.013 | 0.014 | 0.905 | 0.866 | 0.346 | 2.503 |
| 14 | {11000}+{01001,11010,01110,01101}+{01111} | red/red | 0.013 | 0.020 | 0.656 | 0.974 | 0.160 | 6.087 |
| 13 | {11000}+{01001,11010,01101,11111}+{01111} | red/red | 0.013 | 0.020 | 0.656 | 0.974 | 0.160 | 6.087 |
| 12 | {01000}+{11000,01100,01010,01001}+{01110} | red/red | 0.024 | 0.015 | 1.584 | 0.783 | 0.433 | 1.808 |
| 11 | {11010}+{11000,01001,01101,11111}+{11101} | red/red | 0.059 | 0.017 | 3.428 | 0.905 | 0.318 | 2.846 |
| 10 | {11010}+{11000,01001,11011,11111}+{11101} | red/red | 0.059 | 0.017 | 3.428 | 0.905 | 0.318 | 2.846 |
| 9 | {11010}+{11000,01101,11110,11111}+{11101} | red/red | 0.059 | 0.017 | 3.428 | 0.905 | 0.318 | 2.846 |
| 8 | {01010}+{01001,11010,01110,01011}+{01111} | red/red | 0.067 | 0.018 | 3.722 | 0.323 | 0.186 | 1.737 |
| 7 | {01110}+{01001,11010,01101,01111}+{11111} | red/red | 0.075 | 0.022 | 3.483 | 0.841 | 0.136 | 6.184 |
| 6 | {01001}+{11010,01110,01101,01111}+{11111} | red/red | 0.075 | 0.022 | 3.483 | 0.841 | 0.136 | 6.184 |
| 5 | {11011}+{01001,11010,01011,11111}+{01111} | red/red | 0.081 | 0.000 | 1235.241 | 0.126 | 0.996 | 0.127 |
| 4 | {11000}+{01010,01001,11010,01110}+{01011} | red/blue | 0.170 | 0.030 | 5.666 | 0.718 | 0.026 | 27.722 |
| 3 | {11000}+{01001,11010,11011,11111}+{01011} | red/blue | 0.170 | 0.030 | 5.666 | 0.718 | 0.026 | 27.722 |
| 2 | {01101}+{01001,11010,01110,01111}+{01011} | red/red | 0.192 | 0.013 | 15.097 | 0.533 | 0.362 | 1.472 |
| 1 | {01101}+{01001,11010,01111,11111}+{01011} | red/red | 0.192 | 0.013 | 15.097 | 0.533 | 0.362 | 1.472 |

**Table S6:** Significant 4-dimensional interactions, which cannot be seen in lower dimensions, cf. (Fig. S20). The value *p* ↑ refers to the *p*-value of the 4-dimensional bipyramid in question whereas *p* ↓ is the *p*-value of its ridge intersected with the ∩ - face, cf. (Fig. S20c) for the Gould data.

| Data | significant bipyramid | ∩-face | $p \uparrow$ | $p \downarrow$ |
| --- | --- | --- | --- | --- |
| Eble | - | - | - | - |
| Gould |  |  |  |  |
| **0** | {00010} + {00000, 10010, 00011, 11011} + {10001} | **01* | 0.041 | 0.270 |
|  |  | *00** | 0.041 | 0.149 |
|  | {10010} + {00000, 11000, 10001, 11011} + {01001} | 1*0** | 0.041 | 0.076 |
|  |  | **00* | 0.041 | 0.063 |
| Khan |  |  |  |  |
| 0**** | {00010} + {00000, 01001, 00101, 00011} + {00001} | 0***1 | 0.009 | 0.052 |

**Table S7:** Bacterial species considered in the two microbiome data sets.

|  | Gould data set | Eble data set |
| --- | --- | --- |
| Species 1 | *L. plantarum* | *L. plantarum* |
| Species 2 | *L. brevis* | *L. brevis* |
| Species 3 | *A. pasteurianus* | *A. cerevisiae* |
| Species 4 | *A. tropicalis* | *A. malorum* |
| Species 5 | *A. orientalis* | *A. orientalis* |

**Table S8:** Parallel analysis Eble 0**** → 1****, non-critical red/red-case omitted.

| No. | bipyramid<sub>s</sub> | type | $e_o$ | $e_p$ | $e_o/e_p$ | $p_o$ | $p_p$ | $p_o/p_p$ |
| --- | --- | --- | --- | --- | --- | --- | --- | --- |
| 23 | {00001}+{00000,01001,01011,00111}+{01111} | red/red | 0.001 | 0.012 | 0.066 | 0.953 | 0.390 | 2.444 |
| 22 | {00001}+{00000,01001,01101,00111}+{01111} | red/red | 0.001 | 0.012 | 0.066 | 0.953 | 0.390 | 2.444 |
| 21 | {01110}+{00000,00110,01011,01111}+{00111} | red/blue | 0.001 | 0.025 | 0.041 | 0.923 | 0.038 | 24.226 |
| 20 | {01110}+{00000,01100,00110,01111}+{00111} | red/blue | 0.001 | 0.035 | 0.041 | 0.923 | 0.038 | 24.226 |
| 19 | {00110}+{00000,01100,00111,01111}+{01101} | red/red | 0.002 | 0.012 | 0.201 | 0.827 | 0.303 | 2.729 |
| 18 | {00110}+{00000,01100,00101,00111}+{01101} | red/red | 0.003 | 0.014 | 0.201 | 0.827 | 0.303 | 2.729 |
| 17 | {01110}+{00000,01000,01100,01111}+{01101} | red/red | 0.003 | 0.013 | 0.264 | 0.742 | 0.251 | 2.956 |
| 16 | {00110}+{00000,00010,01011,00111}+{00011} | red/red | 0.004 | 0.003 | 1.568 | 0.755 | 0.843 | 0.896 |
| 15 | {00010}+{00000,01010,00110,01011}+{01110} | red/red | 0.007 | 0.010 | 0.748 | 0.606 | 0.488 | 1.242 |
| 14 | {01010}+{00000,00010,00110,01011}+{00111} | red/red | 0.008 | 0.005 | 1.583 | 0.443 | 0.639 | 0.693 |
| 13 | {01010}+{00000,00110,01110,01011}+{01111} | red/red | 0.009 | 0.024 | 0.359 | 0.475 | 0.062 | 7.686 |
| 12 | {01010}+{00000,01000,01110,01011}+{01111} | red/red | 0.009 | 0.024 | 0.359 | 0.475 | 0.062 | 7.686 |
| 11 | {00100}+{00000,01100,00110,00101}+{00111} | red/red | 0.009 | 0.018 | 0.498 | 0.533 | 0.269 | 1.981 |
| 10 | {01001}+{00000,00001,01101,00111}+{00101} | red/red | 0.014 | 0.014 | 1.018 | 0.288 | 0.313 | 0.920 |
| 9 | {00101}+{00000,01100,01101,00111}+{01111} | red/red | 0.015 | 0.026 | 0.584 | 0.228 | 0.062 | 3.695 |
|  | {00101}+{00000,01100,00110,00111}+{01111} | red/blue | 0.018 | 0.040 | 0.446 | 0.321 | 0.035 | 9.119 |
| 8 | {01101}+{00000,01001,00111,01111}+{01011} | red/red | 0.019 | 0.003 | 6.623 | 0.068 | 0.800 | 0.085 |
| 7 | {01101}+{00000,01000,01001,01111}+{01011} | red/red | 0.019 | 0.003 | 6.623 | 0.068 | 0.800 | 0.085 |
| 6 | {01001}+{00000,00001,01011,00111}+{00011} | red/red | 0.019 | 0.005 | 3.571 | 0.153 | 0.689 | 0.222 |
| 5 | {01000}+{00000,01010,01110,01011}+{00110} | red/red | 0.020 | 0.011 | 1.750 | 0.169 | 0.443 | 0.381 |
| 4 | {01000}+{00000,01100,01110,01111}+{00110} | red/red | 0.020 | 0.011 | 1.750 | 0.169 | 0.443 | 0.381 |
| 3 | {01000}+{00000,01001,01011,01111}+{00111} | red/red | 0.021 | 0.013 | 1.535 | 0.140 | 0.339 | 0.413 |
| 2 | {01100}+{00000,01000,01101,01111}+{01001} | blue/blue | 0.045 | 0.037 | 1.215 | 0.000 | 0.003 | 0.176 |
| 1 | {01001}+{00000,01000,01011,01111}+{01110} | blue/red | 0.048 | 0.024 | 1.993 | 0.000 | 0.056 | 0.002 |
|  | {01100}+{00000,01000,01110,01111}+{01011} | blue/blue | 0.064 | 0.034 | 1.855 | 0.000 | 0.005 | 0.000 |
|  | {00010}+{00000,00011,01011,00111}+{00001} | blue/blue | 0.065 | 0.043 | 1.518 | 0.000 | 0.001 | 0.001 |
|  | {01100}+{00000,01101,00111,01111}+{01001} | blue/red | 0.066 | 0.024 | 2.775 | 0.000 | 0.105 | 0.000 |
|  | {00001}+{00000,00101,01101,00111}+{01100} | blue/blue | 0.066 | 0.033 | 1.989 | 0.000 | 0.009 | 0.000 |
|  | {01001}+{00000,01011,00111,01111}+{00110} | blue/blue | 0.068 | 0.036 | 1.917 | 0.000 | 0.007 | 0.000 |
|  | {01100}+{00000,00110,01110,01111}+{01011} | blue/blue | 0.083 | 0.045 | 1.829 | 0.000 | 0.002 | 0.000 |
|  | {01100}+{00000,00110,00111,01111}+{01011} | blue/red | 0.084 | 0.021 | 4.035 | 0.000 | 0.210 | 0.000 |

**Table S9:** Regressions over {0001}+{0000,1001,1011,0111}+{1111} for normalized lifespan data for Eble 0**** and Eble 1****.

|  | Coefficient | Std. error | *t*-statistic | *p*-value |
| --- | --- | --- | --- | --- |
| $\beta_0$ | 0 | 0 | nan | nan |
| $x_1$ | -0.0270 | 0.009 | -2.987 | 0.003 |
| $x_2$ | -0.0149 | 0.012 | -1.246 | 0.213 |
| $x_3$ | -0.0156 | 0.012 | -1.306 | 0.192 |
| $x_4$ | 0.2039 | 0.008 | 26.022 | 0.000 |
| $\beta_0$ | 0.2320 | 0.005 | 44.642 | 0.000 |
| $x_1$ | 0.0310 | 0.005 | 5.957 | 0.000 |
| $x_2$ | 0.0610 | 0.007 | 8.874 | 0.000 |
| $x_3$ | -0.0185 | 0.007 | -2.692 | 0.007 |
| $x_4$ | -0.0861 | 0.007 | -12.518 | 0.000 |

## Notes

https://github.com/holgereble/EpistaticFiltration

### Competing Interest Statement

The authors have declared no competing interest.

